# Early establishment of small intestine neuroendocrine tumors

**DOI:** 10.1101/2025.05.16.654489

**Authors:** Alice Schiller, Anna Hultmark, Markus Lindberg, Andreas Socratous, Tom Luijts, Jimmy Van den Eynden, Anna-Karin Elf, Rauni Rossi Norrlund, Yvonne Arvidsson, Erik Elias, Erik Larsson

**Author notes:** Correspondence to Erik Larsson,. These authors contributed equally (Anna Hultmark, Markus Lindberg, Andreas Socratous, Erik Elias, Erik Larsson).

## Abstract

Small intestine neuroendocrine tumor (SI-NET) is normally diagnosed late in life and has several unusual properties, including low proliferation rate, low mutation burden, lack of driver mutations, and frequent multifocality in the form of polyclonal tumor clusters. This sets SI-NET aside from other adult cancers and raises questions about the timeline of initiation and progression. Here, we investigated the evolutionary history of multifocal and unifocal SI-NET using whole genome sequencing data, obtaining timing information on primary tumors, metastases and key genetic events. Despite the late onset, the results indicated that major genetic alterations and the establishment of advanced metastatic tumor cell clones can often be traced back to childhood or adolescence in SINET. This was validated by re-examination of archival CT/MR scans, allowing longitudinal tracking of individual tumors up to 12 years prior. Metastases were detected at a high degree of consistency in the historical imaging data and estimated growth rates suggested several additional decades of tumor expansion. Collectively, our data support that slow growth of advanced lesions over half a century or more may precede SI-NET diagnosis in late adulthood.

## INTRODUCTION

Small intestine neuroendocrine tumor (SI-NET), the most common cancer of the small intestine, has several distinct properties, such as a low proliferative capacity and a remarkably low mutational burden^1-3^. Unlike many other adult cancers, and despite often being metastatic at diagnosis^4,5^, SI-NET essentially lacks identifiable somatic driver mutations that can explain their initiation and spread. Moreover, approximately 50% of SI-NET cases display multifocality, where several clonally independent tumors grow in proximity^6,7^. As SI-NET is commonly diagnosed at a late stage with advanced metastatic spread there is a need for new systemic treatment options, but development is hindered by the poor understanding of the underlying mechanisms driving tumor initiation and progression.

The late clinical presentation of SI-NET, typically in patients over the age of 60, has historically led to the assumption that tumorigenesis is a relatively late-onset event. However, some of the biological characteristics of SI-NET share parallels with pediatric cancers, which are known for their low mutation burden and relative absence of canonical driver mutations, relying instead on copy-number alterations, non-genetic mechanisms or early developmental events to fuel tumorigenesis^8^. This comparison motivates a closer investigation into the evolutionary timeline of SI-NET.

In this study, we analyze whole-genome sequencing (WGS) data and re-examine archival clinical imaging data to track the evolutionary history of multifocal and unifocal SI-NETs. Our findings indicate that key molecular events and the establishment of advanced tumor cell clones can often be traced back to childhood or adolescence, and that clinical detection may thus commonly be preceded by more than 50 years of tumor growth.

## RESULTS

### Overview of molecular alterations in 74 SI-NETs from 16 patients

To gain insight into evolutionary histories in SI-NET, we first combined whole genome sequencing (WGS) data from 11 patients with multifocal tumours^6^ with additional WGS of five unifocal cases, for a final cohort encompassing 47 primary tumors, 27 metastases, 7 adjacent normal tissue samples and 16 normal blood samples (**Fig. 1a-b**). The age at surgery averaged 70.5 years across the 16 SI-NET patients and was similar in multi- and unifocal cases (71.5 vs. 68.2 years; **Fig. 1c; Supplementary Data 1**). The data was processed using a standardized pipeline that included strict somatic single nucleotide variant (SNV) calling and filtering to minimize false positives (see **Methods**). Mutations were nearly absent in the normal tissue samples as expected (8-15 SNVs genome-wide), supporting that mutation calls were highly specific (**Supplementary Fig. 1**; **Supplementary Data 1**).

**Figure 1.**
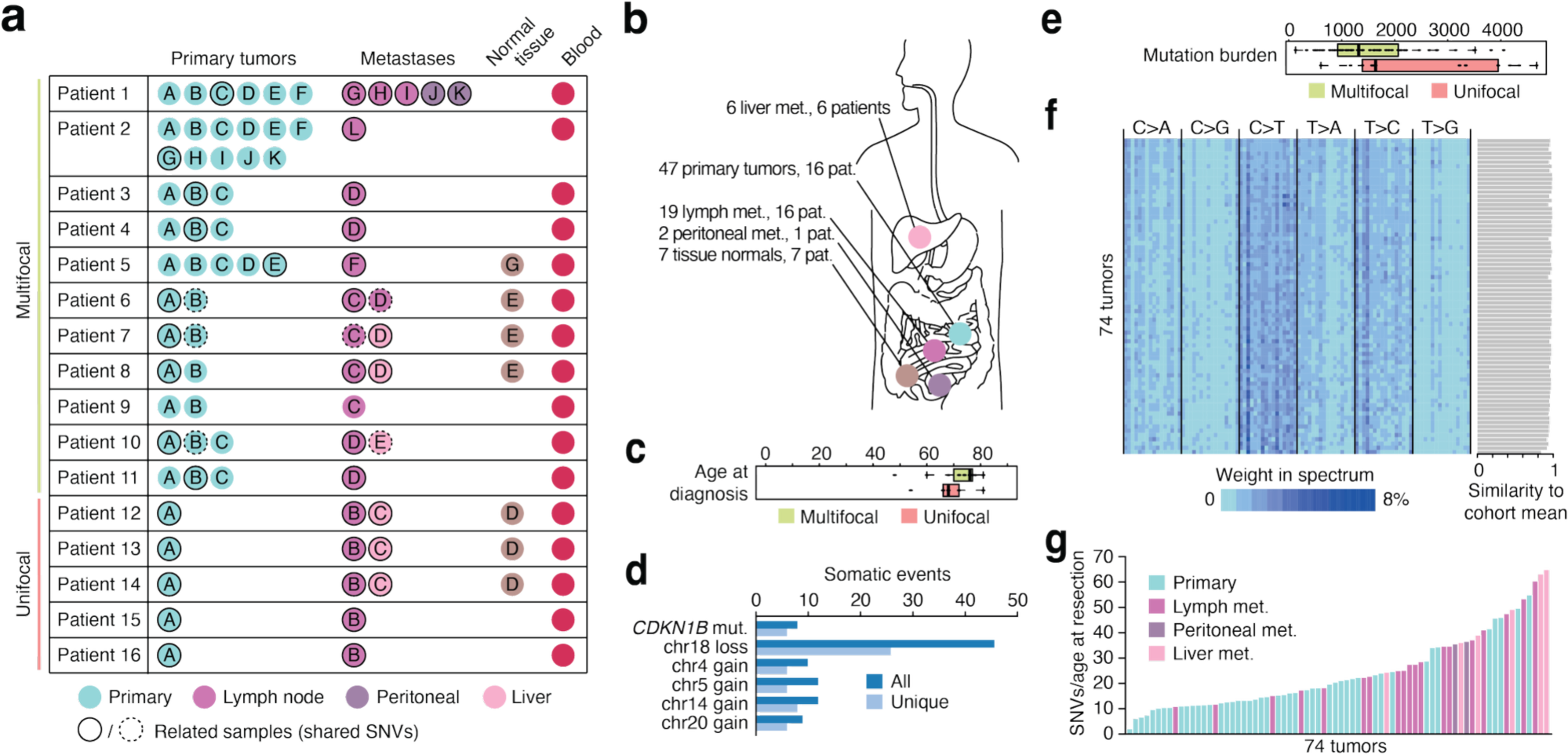
Cohort composition and key mutational characteristics. (**a**) Overview of whole-genome-sequenced samples included in the study, each represented by a filled circle. Evolutionarily related primary tumors and metastases, identified based on overlapping somatic SNVs, are indicated by black circles. Where applicable, dashed black circles are used to indicate a second, independent, evolutionary clade. Primary (intestinal) tumors were unrelated as indicated by lack of shared somatic mutations. (**b**) Anatomical overview of sample origins. (**c**) Age at surgery/diagnosis in multi- and unifocal cases. (**d**) Key recurrent somatic events (detailed overview in **Supplementary Fig. 1**). (**e**) Somatic SNV burden in multi- and unifocal cases. (**f**) Heatmap showing that the overall trinucleotide substitution spectrum is similar across tumors, with cosine similarity to the cohort mean indicated for each sample. Samples were sorted by mutation burden (low to high). (**g**) SNV counts normalized to age at resection in each sample, which will underestimate the yearly mutation accumulation rate but can approach it in case of late clonal expansions.

Multifocal lesions were clonally independent as indicated by a general lack of shared somatic SNVs, while metastases could generally (15/16 patients) be confidently traced back to specific seeding primary tumors based on numerous overlapping SNVs, confirming earlier analyses (**Supplementary Fig. 1**; **Supplementary Note 1**)^6,7^. Likewise, the pattern of recurrent somatic alterations confirmed earlier data^1,9^, with loss of chromosome 18 being the most frequent copy number event, seen in 46 samples (26 unique events), followed by gains of chromosomes 4, 5, 14 and 20 (**Fig. 1d**; **Supplementary Fig. 1**). Notably, two of the unifocal cases exhibited a distinctive copy number instability pattern with smaller (∼3-10 Mb) focal amplifications scattered throughout the genome (**Supplementary Fig. 2**).

Disrupting coding variants in *CDKN1B*, the most common putative driver mutation^1^, were present in 8 tumors (6 unique events; **Fig. 1d**) and were sometimes found in seeding primary lesions while being absent in the corresponding metastases, as previously reported^6^ (**Supplementary Fig. 1**; potential driver mutations listed in **Supplementary Data 2**). Additionally, a somatic frameshift variant in *TNRC6B*, previously pointed out as a potential driver^7^, was present in one non-seeding multifocal intestinal tumor (**Supplementary Fig. 1**).

### Mutagenesis in SI-NET resembles normal cells

SNV counts ranged from 148 to 4,660 genome-wide (0.05-1.50 per Mb), confirming a low burden^1,6,7^ (**Supplementary Fig. 1**). A comparison with available data from other cancer types^10^ further underscored that SI-NET exhibits relatively low SNV counts despite the generally advanced age at diagnosis (**Supplementary Fig. 3**). While the median count was somewhat higher in unifocal compared to multifocal cases (1,645 vs. 1,329, respectively; *P* = 0.046, Wilcoxon rank sum test; **Fig. 1e**), a higher proportion of the unifocal samples were metastases with overall higher burdens, and there was no significant difference when considering intestinal tumors or metastases alone (*P* = 0.30 and *P* = 0.18, respectively).

Consistent with the low mutation burdens, SNV trinucleotide mutation spectra were highly similar across tumors, supporting that no individual samples were notably exposed to distinct or atypical mutagenic processes (**Fig. 1f**). Deconvolution of contributions from COSMIC^11^ signatures, while complicated by the relatively low mutation counts, supported that SBS5 (a clock-like process) accounted for the majority of mutations in nearly all samples (**Supplementary Fig. 4**). Additional, smaller, contributions were consistently observed from SBS1 (deamination of 5-methylcytosine), SBS40 (clock-like), SBS3 (homologous recombination deficiency) and SBS8 (unknown).

Notably, SBS5, SBS40, SBS3 and SBS8 are all relatively featureless and similar, and have previously been described as “flat” signatures^12^ (**Supplementary Fig. 5**). Their relative contributions varied considerably between bootstrap iterations in individual samples, indicative of low solution stability, as well as between analysis tools (deconstructSigs^13^ and SigProfiler^14^; **Supplementary Fig. 4**). As such, their exact individual contributions cannot be robustly resolved and should be interpreted with caution. Furthermore, there were no convincing differences in signatures in early (truncal shared by related tumors and metastases) compared to later (non-truncal) SNVs (**Supplementary Fig. 6**). Together, the uniformity in trinucleotide spectra, along with the dominance and consistently high proportion of clock-like/aging-related mutations across samples, supports that mutagenesis in SI-NET is largely normal and stable over time.

Next, we related the SNV burden in each sample to patient age at resection, which will consistently underestimate the true yearly mutation accumulation rate but can approximate it in case of a late clonal expansion. We found that the SNV/age ratio peaked at 64.7 SNVs/year, with the highest values observed in two distant metastases (liver), potentially reflecting relatively late expansions (**Fig. 1g**). In conclusion, while some minor additional exposures cannot be excluded, mutagenesis and mutation rates in SI-NET appear to largely mirror normal cells, with ileum and colon epithelial cells acquiring an estimated 42 and 47 SNVs/year, respectively, for comparison^15,16^.

### Genetic timelines in SI-NET support early initiation

We next sought to estimate the timeline of major genetic events in SI-NET. In particular, a timepoint for the most recent common ancestor (MRCA) tumor cell clone preceding both the primary tumor and related metastases can be estimated based on the number of shared (truncal) SNVs in phylogenetic analyses. To ensure sufficient sensitivity for detecting shared and private variants, we first examined the variant allele read count distributions across all primary–metastasis pairs. Most fell well safely within the detectable range, suggesting limited dropouts (**Supplementary Fig. 7**). This was supported by results from patient 1, where the same set of early truncal variants was consistently recalled across six related tumors with limited variation (909–937 SNVs; **Supplementary Note 1**).

We found that, despite their high age at surgery, the primary/metastasis MRCA clones were often represented by surprisingly few mutations (average 992, ranging from 148 to 2,969) and thus traceable back to early timepoints in life, in both multifocal and unifocal cases (**Fig. 2a** and **Fig. 2b**, respectively; detailed case analyses in **Supplementary Note 1**). Based on an assumption of 40-80 SNVs accumulated per year, the MRCA clones would have been established at an average age of 12.4-24.8 years across the cohort (earliest case 1.9-3.7 years), with higher mutation rate assumptions leading to lower age estimates (**Fig. 2c**; **Supplementary Data 3**).

**Figure 2.**
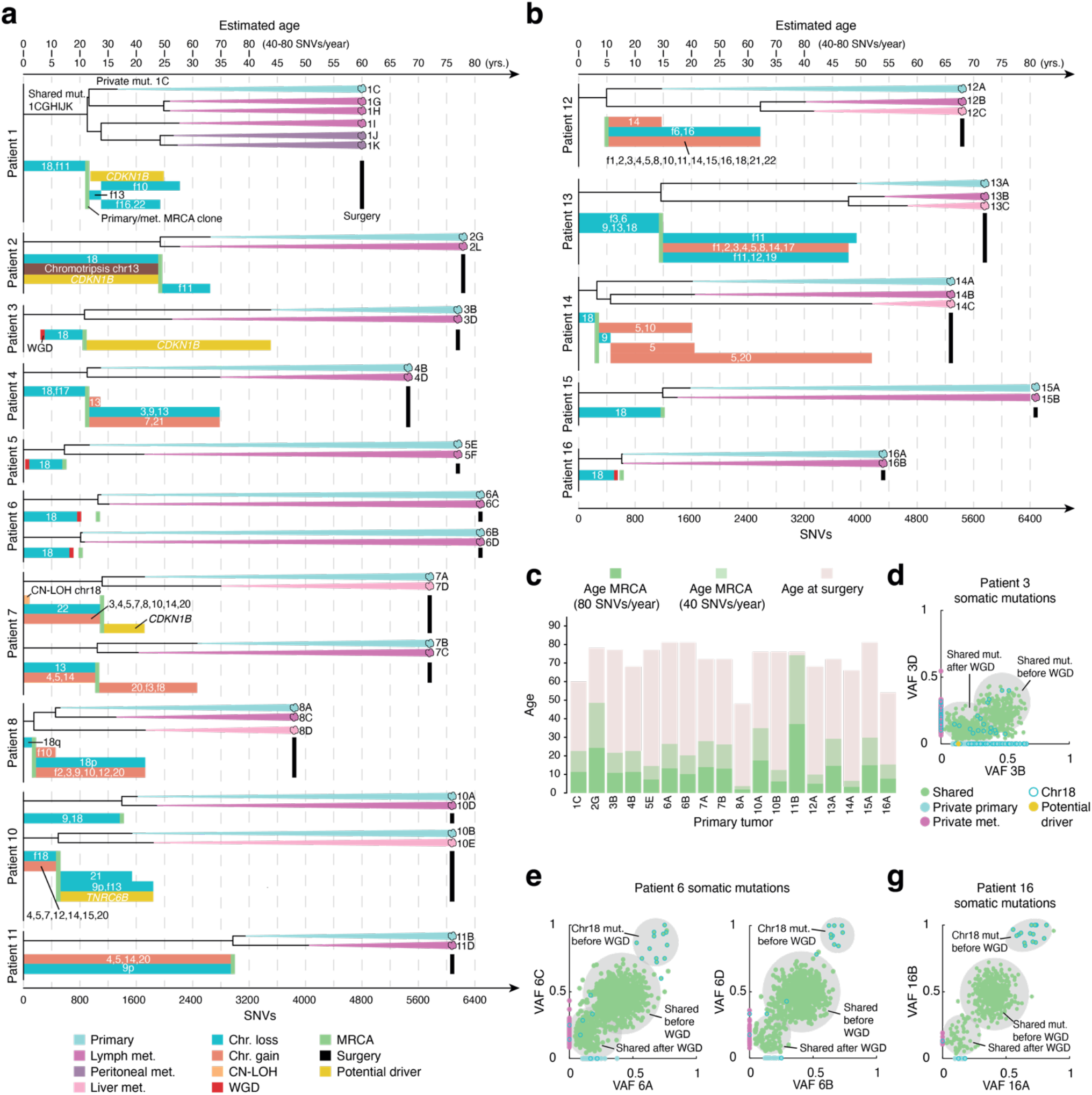
Genetic timelines of evolutionarily related SI-NET primary tumors and metastases. (**a**) Maximum parsimony phylogenetic trees of primary-metastasis evolutionary clades in multifocal SI-NET patients, based on somatic SNVs. The number of SNVs in each branch in indicated by the scale at the bottom. Approximate corresponding timepoints, assuming a mutation rate of 40 or 80 SNVs/year, are given at the top. The mutation burden and approximate age of the most recent common ancestor (MRCA) clone of primary tumors and metastases is given by the length of the trunk in each tree (green bars). The age at resection/surgery is indicated for each sample (black bars). Timepoints or intervals for other major genetic events (e.g., chr18 deletion) are indicated below each tree. Patient 9 (not shown) lacked a primary-metastasis pair. “f” indicates a focal or smaller copy number event, while “p” and “q” indicate an arm-level events. (**b**) Same as panel **a** but for unifocal SI-NET cases. (**c**) Overview of the estimated age of all tumor/metastasis MRCA clones, based on an assumption of a yearly accumulation of either 40 or 80 SNVs/year (light or dark green, respectively), shown together with the age at surgery/diagnosis. (**d**-**g**) Detailed view of SNVs and their variant allele frequencies (VAF) in select tumor-metastasis pairs.

Similarly, total mutation burdens were in several samples low enough to support that considerable time had passed between the last major clonal expansion and tumor resection. For example, the metastatic primary tumors in patients 4, 5, 6 and 16 had SNV counts ranging from 622 to 1,098, with most variants being clonal and with high sample purity as indicated variant allele frequency (VAF) distributions, equivalent to a chronological age of 15.6 to 27.5 years at 40 SNVs/year or 7.8 to 13.7 years at 80 SNVs/year (**Fig. 2a-b**).

While metastases typically had higher burdens, consistent with later clonal expansions while still not excluding the possibility of early seeding, a few had SNV counts low enough to support that metastasis had occurred already in adolescence (627 to 1,213 for metastases in patients 6 and 16, equivalent to 15.7 to 30.3 years at 40 SNVs/years or 7.8 to 15.2 years at 80 SNVs/year; **Fig. 2a-b**). While the timepoints are approximate with a large margin of error, these findings nonetheless suggest that not only precancerous lesions, but also advanced tumors and metastases commonly date back many decades prior to diagnosis, consistent with the slow-growing nature of SI-NET.

### Chromosome 18 deletion and whole genome doubling occurs early in life

We next estimated timepoints for key somatic genetic events SI-NET, aided by inheritance patterns in the established phylogenetic relationships (detailed in **Supplementary Note 1**). Deletion of chromosome 18, the main recurrent event, was found in 14/18 metastatic primary tumors, and was always present in the corresponding metastases (**Supplementary Fig. 1**). Furthermore, phasing based on germline single nucleotide polymorphisms (**Supplementary Note 1**) showed that the same chromosome homolog was always lost in these cases, despite both paternal and maternal homolog losses otherwise being observed within the same patients (*P* = 2.4×10^−4^; binomial test based on 12 metastatic primary tumors with monoallelic loss). This rules out late convergent events, allowing the conclusion that chromosome 18 loss consistently emerges early, between embryogenesis and establishment of primary/metastasis MRCA clones (**Fig. 2a-b**). While other events could sometimes also be truncal (present in primary and metastasis), many would often arise later, including focal copy number changes and *CDKN1B* mutations (**Fig. 2a-b**; further supported by results from MutationTimer^17^, **Supplementary Note 2**). These findings support that early chromosome 18 deletions may play a key role in SI-NET development.

Whole genome doubling (WGD), while relatively frequent in human cancers overall^18^, is not well-established in SI-NET. In four patients and ten tumors, SNV VAF distributions and their somatic inheritance patterns supported early truncal WGD events, before tumor/metastasis MRCA clones but typically after chromosome 18 deletion (**Fig. 2d-g** and **Fig. 2a-b**; **Supplementary Note 1**). In Patient 3, diagnosed at age 76, our analysis supported that WGD likely occurred at an age of only a few years, before chromosome 18 deletion (**Supplementary Data 3**). WGD thus has a possible functional role in SI-NET and can arise in early childhood even in patients diagnosed at old age.

### Archival CT scans from two of the patients support early metastatic seeding

SI-NET lesions can be radiologically difficult to distinguish from normal findings in computer tomography (CT) or magnetic resonance (MR) examinations and are sometimes only recognized retrospectively after functional imaging. We therefore hypothesized that archived CT or MR scans might help corroborate the findings from our genomic analyses and reviewed archival imaging data for all 16 patients. The digital archive extended back about 15 years, limited by historical storage practices and the less frequent use of CT in earlier diagnostics. In three cases (patients 6, 11 and 16), we identified abdominal CT scans performed several years prior to SI-NET diagnosis, two of which revealed previously undetected tumors identifiable in the light of findings at diagnosis.

Strikingly, in patient 16, we found that a lymph node metastasis present in the pelvic region at surgery (age 54 years) could be identified as a macroscopic lesion (22×17×27 mm; 5.2 cc) upon re-examination of a CT scan taken 11 years earlier (**Fig. 3a**; **Supplementary Data 4**). In this case, the genetic analysis had revealed that chromosome 18 deletion, WGD and tumor/metastasis evolutionary divergence had likely occurred in childhood (**Fig. 2b, Fig. 3a**). Additional CT data collected four years before diagnosis, as well CT and MR data obtained at diagnosis, supported approximately exponential growth during the 11-year time span (**Fig. 3a**). Interpolation of exponential tumor growth backwards in time by regression analysis was consistent with metastatic seeding having occurred at a much earlier age, likely already in childhood (**Fig. 3a**).

**Figure 3.**
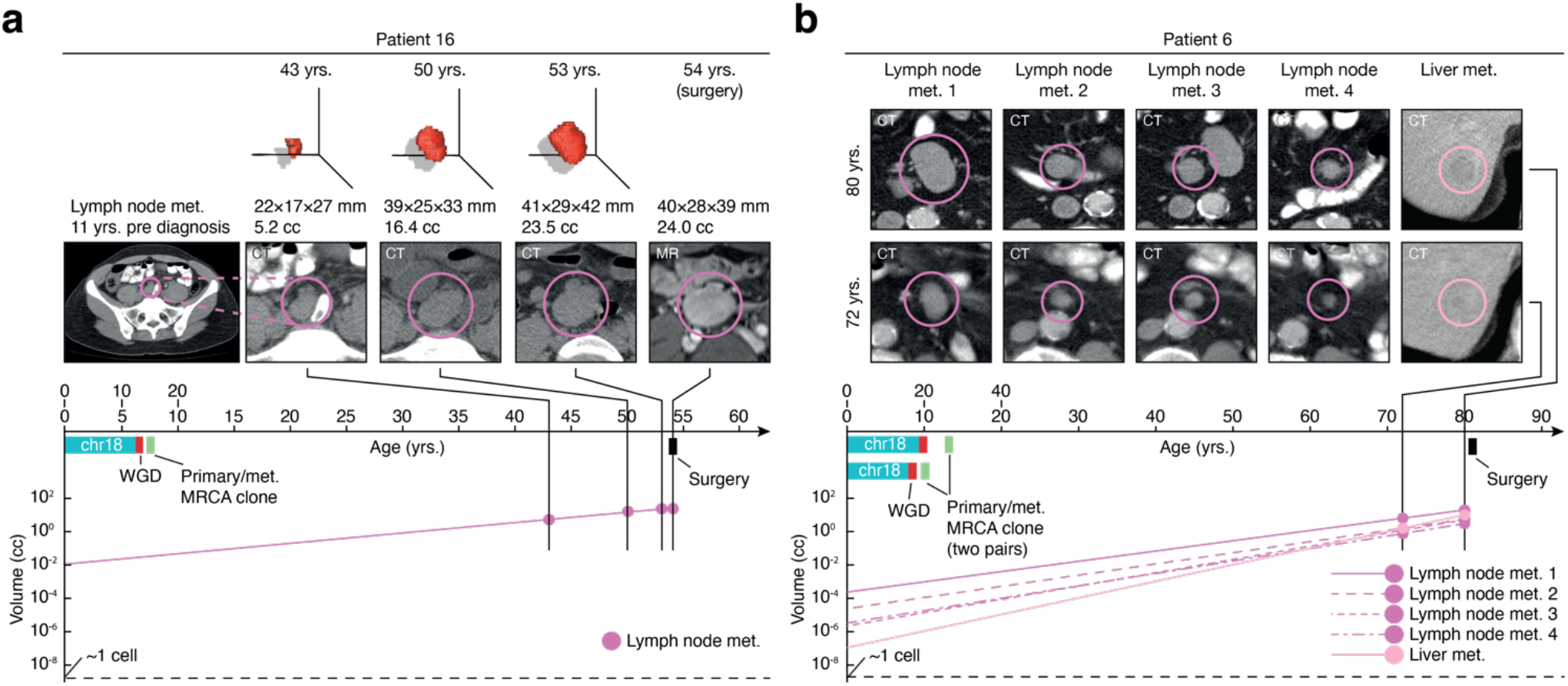
Archival tomography scans from two of the patients confirms early metastasis. (**a**) In patient 16, a lymph node metastasis found at diagnosis (age 54 years) was identified as a prominent macroscopic lesion (22×17×27 mm) in an archived CT scan taken 11 years earlier. Regression together with late and intermediate timepoints support approximately exponential growth during the period and is compatible with metastatic seeding having occurred at a much earlier age (linear regression line shown on a logarithmic volume scale). The dotted line indicates the approximate volume of a single cell (2×10^−9^ cc)^19^. (**b**) Similar results from patient 6, where four lymph node metastases and a liver metastasis could all be readily identified as macroscopic lesions in CT data 9 years prior to surgery. The plots show volume estimates with regression lines (exponential model) supporting that metastasis events were considerable older, possibly occurring even in childhood. Key genetic events are indicated for both patients, together with age estimates assuming 40 or 80 SNVs/year: early chr18, early WGD and early tumor/metastasis MRCA clones. MR, magnetic resonance imaging.

Similarly, in patient 6, where the genetic data had also pointed to early-in-life chromosome 18 deletion, WGD and MRCA clones for two sequenced tumor/metastasis pairs, we found that four lymph node metastases and a liver metastasis present at surgery were all readily identifiable as macroscopic lesions in CT data acquired 9 years earlier (**Fig. 3b**). Metastases must thus have been present at considerably earlier timepoints, and under the assumption of exponential tumor growth, monoclonal metastatic seeding would have occurred in childhood (**Fig. 3b**).

### Additional archival CT/MR data further supports early growth and metastasis in SI-NET

To further investigate the timeline of tumor initiation and growth in SI-NET, we systematically reviewed archival imaging data for 102 additional SI-NET patients that were not part of the original WGS cohort, identifying 13 cases with relevant preoperative scans. For many patients, multiple CT or MR scans had been performed at different timepoints, and in nearly half of the cases (6/13) the earliest scans were done at least 10 years before diagnosis (**Fig. 4a**; **Supplementary Data 4**).

**Figure 4.**
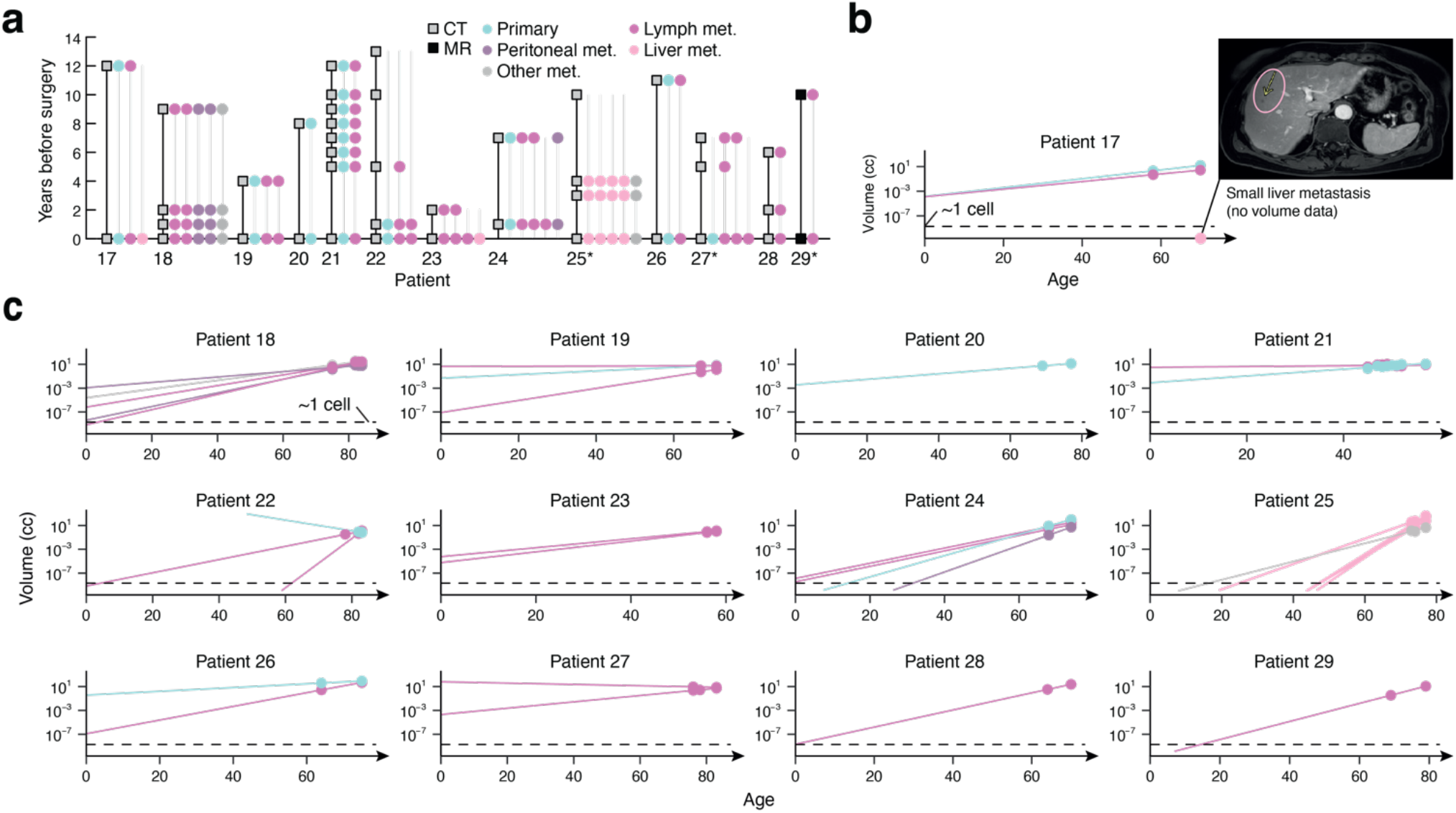
Early-in-life metastatic SI-NET supported by archival tomography data from 13 additional patients. (**a**) Overview of patients and timepoints for CT/MR scans and detections of tumors. Grey lines indicate the timespan where informative data was available for each tumor. *In patients 25 and 27, failed detection of a pulmonary metastasis (grey) 10 years before diagnosis and a lymph node metastasis 6 years before diagnosis, respectively, was due to missing relevant imaging data. In patient 29, CT was also performed at diagnosis, but for consistency, only MR data was considered. (**b-c**) Volume estimates for individual lesions at different timepoints (logarithmic volume scale; regression lines indicate growth under an exponential model). The dotted line indicates the approximate volume of a single cell (2×10^−9^ cc)^19^.

In all 13 cases, SI-NET tumors identifiable by functional imaging at diagnosis could be identified also at earlier timepoints (**Fig. 4a**; **Supplementary Data 4**). Notably, with only two exceptions (patients 22 and 25), all scans were tumor positive, and not only metastases but also primary tumors could be traced up to 12 years back in time (**Fig. 4a**). The earlier detections were thus constrained primarily by availability of historical data, supporting that latency periods were often considerably longer than indicated by the timepoints of the oldest available scans. Interpolation of tumor growth backward in time, while based on a simplified exponential growth model of tumor expansion and complicated by uncertain volume estimates for small primary tumors in some cases, was compatible with monoclonal expansion having started already in childhood in most cases (**Fig. 4b-c**).

## DISCUSSION

Our study provides strong support from both genetic and clinical imaging data that the initiation of SI-NET often occurs early in life, with tumorigenesis and even metastasis likely having occurred by adolescence or childhood in many cases. This changes current paradigms framing this disease as one of late adulthood. While there is a growing body of evidence that neoplastic transformation may start earlier than previously appreciated for many adult cancers^17^, the extreme timespan suggested by our results, where even metastatic seeding may plausibly have occurred more than half a century before diagnosis, is unusual yet compatible with the low proliferation rate of SI-NET^20^.

Our findings have implications for early detection and clinical management. A major obstacle to improving outcomes in SI-NET is that diagnosis often occurs at a late stage, when distant metastases are already present and curative treatment by complete surgical resection is no longer feasible. Earlier detection of disease is therefore highly warranted. Our data suggest that SI-NET should be considered in younger patients with unexplained bowel symptoms, and that individuals with a familial history may benefit from screening at an early age.

The genetic analyses can only estimate the timepoints of latest clonal expansion that seeded metastases, which still leaves the possibility that initiation may have occurred even earlier, potentially during embryogenesis. Although this is consistent with prior speculation regarding a developmental origin of multifocal SI-NET^21^, especially given that multifocal SI-NETs are clonally independent^6,7^, the exact timing and mechanism of initiation remains unknown. While our results reinforce a possible role for chromosome 18 deletions, pointing to their consistently early emergence, more work is needed to explore the possibility of non-genetic initiating factors, including early epigenetic reprogramming and microenvironmental influences.

To independently validate our genetic results, we reassessed historical CT and MR data. While cases with earlier scans were understandably limited and may not be representative of all SI-NET patients, a striking proportion (15/16) had detectable tumors many years before diagnosis. Importantly, the radiology data revealed not only the presence of tumors at earlier timepoints but also confirmed the slow growth rate of SI-NETs. While extrapolation beyond observed data points does not allow for precise estimation of lesion size at specific timepoints, and should be interpreted with caution, the results still strongly support that metastatic spread often predated the earliest available scans by many additional decades, in agreement with the genomic data.

Together, our findings redefine the timeline of SI-NET development and raise fundamental questions about early initiating events and the basis for clonally independent multifocal tumors. This study lays the foundation for future mechanistic research and opens new avenues for early detection.

## METHODS

### Patient cohort and sample collection

The study cohort evaluated for inclusion in the present study consists of patients that have underwent surgery for SI-NET at Sahlgrenska University Hospital and provided informed consent. Study participation includes sampling of tumor and normal tissue during surgical procedures and retrieval of clinical information from hospital journals. Participants receive in-person information by a designated research nurse and written information explaining the purpose of the study. In accordance with the ethical permit, consent is obtained in-person and documented in writing by a signed document or a designated entry in the patient’s medical journal. The study was approved by the Central ethical review board in Gothenburg (Dnr. 833-18).

The present sequencing cohort consists of 11 previously sequenced patients with multifocal disease^6^ combined with an additional 5 patients with only one detectable (by palpation during surgery) intestinal tumor (i.e. unifocal disease). Tissue sampling was performed as previously described^6^. In brief, samples from tumor tissue and normal tissue was collected in association with surgery and snap-frozen in liquid nitrogen. A piece of each individual tissue sample was formalin-fixed and paraffin-embedded for additional studies by immunohistochemistry. Collected tissue was evaluated by a pathologist prior to further analysis. Blood was also collected for reference. A diagnosis of well-differentiated neuroendocrine tumor grade 1-2 of the ileum (WHO 2019) was validated according to clinical routine prior to further studies.

### DNA extraction and sequencing

DNA from fresh-frozen biopsies as well as blood from each patient was isolated using the allprep DNA/RNA Mini Kit (Qiagen) according to the manufacturer’s protocol. Libraries were prepared using the TrueSeq Nano kit (Illumina) and were sequenced on an Illumina Novaseq 6000 using 150 bp paired-end reads. The average sequencing depth across all 97 samples considered in the study ranged from 29.5× to 63.8× (**Supplementary Data 1**).

### Read mapping, sequencing depth and somatic variant calling

The reads were aligned to hg38 using bwa mem (v 0.7.17-r1188) with option -M. Deduplication and Base Quality Score Recalibration (BQSR) were performed with MarkDuplicates respectively ApplyBQSR from GATK toolkit (v4.1.5.0). The approximate average sequencing depth was calculated by (number of mapped reads x 150) / (3.1 × 10^9^).

Variant calling was performed with Mutect2 and VarScan^22^. Mutect2 followed by FilterMutectCalls was run with default settings. VarScan somatic was run with default settings followed by VarScan processSomatic used to filter for somatic SNVs and indels.

Mutations detected by Mutect2 and VarScan were considered high confidence calls if there was no variant reads in the blood normal and a minimum coverage at 20 reads in the normal. In Mutect2 one variant read was required (no strand filter) and in VarScan one variant read per strand was required. Mutations detected in the blood normal sample (called as Germline or LOH in VarScan) of at least two patients were removed. Annotation was performed with VEP.

To increase the sensitivity for pairwise shared subclonal variants while minimizing false positive calls, high-confidence mutations detected by both callers (as described above) in one of the samples in a related pair (sharing > 10 variants) were whitelisted to be included in the other sample if detectable even as a low-confidence call (detectable by one caller without any additional filters).

### Mutational signatures

Mutational signatures were assigned to COSMIC^11^ (version 3.2) using SigProfilerAssignment^23^ (version 0.1.4) and deconstructSigs^24^ (version 1.9.0). Both tools were run without considering treatment signatures as defined by SigProfilerAssignment (SBS11, SBS25, SBS31, SBS32, SBS35, SBS86, SBS87, SBS90, SBS99) and deconstructSigs was run with a limit of 4 signatures per sample. 500 replications using random sampling with replacement of mutations were performed for both tools, and average loadings were determined based on these results.

### Phylogenetic reconstruction

Phylogenetic reconstruction was performed with MEGAX (v10.2.6)^25^. SNVs from the filtered calls were used to build maximum parsimony phylogenies with 500 bootstraps.

### Copy number analysis, purity, homolog phasing and whole genome duplication

Purity was determined for all samples using PurBayes^26^ with default settings. CNAs were called using Battenberg^27^ (version 2.2.10) with default settings. One sample (7C) was run using a purity prior (retrieved from PurBayes) since this sample did not converge to a reasonable purity using default settings.

To decide if the same chromosome homolog was altered within a tumor pair/clade, phasing of CNAs based on heterozygous germline SNPs was performed. For this analysis, LOH and germline calls from VarScan somatic with coverage >=20 and 0.25<=VAF<=0.75 were used.

Manual annotation of WGD was performed for shared primary-metastasis pairs based on VAF distribution of shared and private mutations. WGD was annotated when shared mutations at copy number neutral chromosomes followed a bimodal distribution (representing mutations before and after WGD) and the private mutations followed a unimodal distribution (representing low VAF mutations after WGD). Mutations were divided into the two clusters based on k-means clustering (Euclidian distance).

### Timing of genomic events

The number of shared SNVs between tumor pairs or clades (SNV based phylogeny) was used to decide the MRCA timepoints. The chronological age when the MRCA clones were established, or when clonal expansions occurred, was estimated based on an assumption that 40-80 SNVs accumulate yearly.

CNAs and potential driver mutations were timed in relation to the MRCA timepoint depending on if the event was shared between tumor pairs/clades. Timing of CNAs was further supported by chromosome homolog phasing. CNAs with amplitude (Battenberg logR score) > 0.2 were considered. Timing of WGD events was based on the proportion of all primary/metastasis shared mutations on copy number neutral chromosomes having high VAF (before WGD) as determined by k-means clustering (see above).

MutationTimeR^17^ (default settings) was used to provide timing data on copy number gains in all relevant all primary tumor-metastasis pairs.

### Radiology

A cohort of 118 patients who underwent surgery for SI-NET at Sahlgrenska University Hospital between 2011 and 2022, including the 16 who had undergone genomic sequencing, was selected. The medical records were scrutinized for prior radiological examinations, acquired before diagnosis, that could potentially have detected abdominal tumors. In 17 patients, including 3 of the sequenced patients, older abdominal imaging examinations performed ≥365 days before surgery were identified.

One of the 17 patients was not informative, since only an older CT scan of the upper abdomen was available, which did not include the lower abdomen and pelvis where a small bowel tumor and a mesenteric metastasis were detected on a later preoperative scan. Among the remaining 16 patients with relevant older scans, 15 demonstrated identifiable lesions, two of which were also part of the sequenced patient cohort.

A total of 91 imaging examinations were reviewed and used for quantitative analysis, comprising 79 CT scans, 7 MR scans, and 5 PET/CT scans. The primary focus was on preoperative imaging; all available relevant examinations performed before surgery were included. In selected cases, postoperative studies were also reviewed. Of the 91 examinations, 60 were preoperative and 31 were postoperative. Among the preoperative examinations, 32 were performed ≥365 days prior to surgery, while 28 were acquired within 365 days of surgery.

In total, 60 lesions were identified and followed longitudinally. These included 35 mesenteric metastases, 13 hepatic metastases, 10 small bowel tumors, 1 retroperitoneal metastasis and 1 pulmonary metastasis. The primary emphasis was placed on lesions visible on older preoperative imaging. All identifiable lesions from these studies were included. Additional lesions observed on more recent preoperative studies, and selectively those appearing only on postoperative imaging, were also recorded. Of the 60 lesions, 57 were detectable on at least one preoperative scan, while 3 were only identified postoperatively. Among the 57 preoperative lesions, 39 were visible on imaging performed ≥365 days prior to surgery, while 18 were not. Notably, 9 of these 18 lesions were in patients who had no older imaging available.

Lesion dimensions were measured in three planes: the longest diameter in the axial plane, a perpendicular axial diameter, and a craniocaudal diameter measured in either the sagittal or coronal plane. In some older studies, the slice thickness was too large to allow reliable measurement of the craniocaudal dimension.

Lesion volumes were estimated using a segmentation tool integrated into the PACS (Picture Archiving and Communication System). Small bowel tumors were particularly challenging to quantify, and their volume measurements should be interpreted with caution. Corresponding annotated images with measurement overlays were saved for reference. For visualization and regression, timepoints were rounded to whole years.

## DATA AVAILABILITY

WGS sequencing data that support the findings in the article are in part available under controlled access at the European Genome-Phenome Archive (EGA; https://ega-archive.org), which is hosted by the European Bioinformatics Institute (EBI) and the Centre for Genomic Regulation (CRG), through the primary accession code EGAD00001008831. Access requests should be addressed to Erik Elias or Erik Larsson.

## ACKNOWLEDGEMENTS

The work described here was supported by the Swedish Research Council (E.L.), the Swedish Cancer Society (E.L. and E.E.), the Knut and Alice Wallenberg Foundation (E.L.), grants from the Swedish state under the agreement between the Swedish government and the county councils, the ALF agreement (E.E.), and Kom op tegen Kanker (Stand up to Cancer), the Flemish cancer society (J.V.E.). We wish to thank all the patients who participated in this study.

## COMPETING INTERESTS

None.

## Notes

### Competing Interest Statement

The authors have declared no competing interest.

### Summary of Updates

Title and abstract adjusted to better reflect key results.

