## Supplementary Appendix for "Early establishment of small intestine neuroendocrine tumors"

### Supplementary appendix, Schiller et al.


*Supplementary Figure 1* 2
*Supplementary Figure 2* 3
*Supplementary Figure 3* 4
*Supplementary Figure 4* 5
*Supplementary Figure 5* 6
*Supplementary Figure 6* 7
*Supplementary Figure 7* 8
*Supplementary Note 1* 10
Patient 1 11
Patient 2 12
Patient 3 13
Patient 4 14
Patient 5 15
Patient 6 16
Patient 7 17
Patient 8 19
Patient 9 20
Patient 10 21
Patient 11 22
Patient 12 23
Patient 13 24
Patient 14 25
Patient 15 26
Patient 16 27
*Supplementary Note 2* 28
Patient 7 29
Patient 10 30
Patient 11 31
Patient 14 32
*Supplementary references* 33

### Supplementary Figure 1

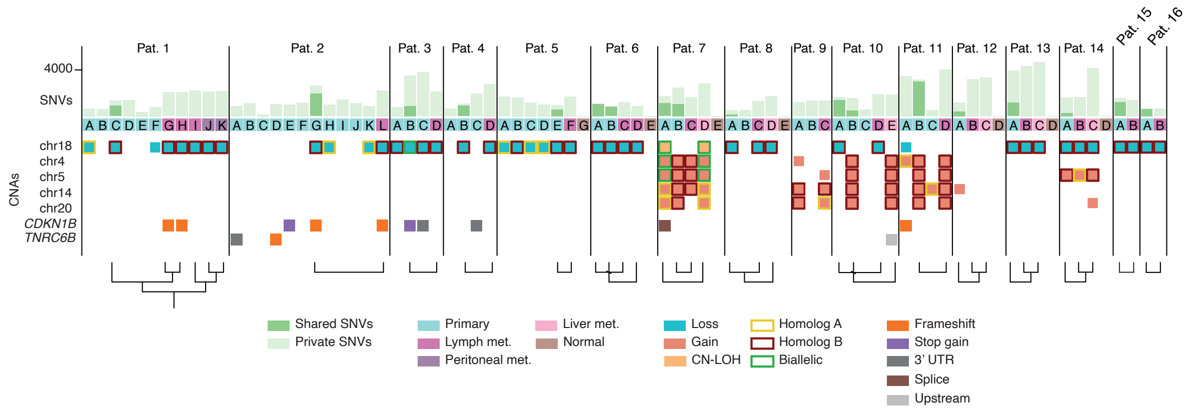


**Detailed overview of somatic genetic alterations for 16 SI-NET patients.** The number of private and shared SNVs are present at the top. The SNV based phylogenetic relationships (further detailed in Supplementary Note 1) are shown at the bottom. Presence of the most common chromosome alterations and mutations in *CDKN1B* and *TNRC6B* are shown. The homolog that was lost/gained is indicated based on the results from phasing of CNAs based on heterozygous germline SNPs (Supplementary Note 1).

### Supplementary Figure 2

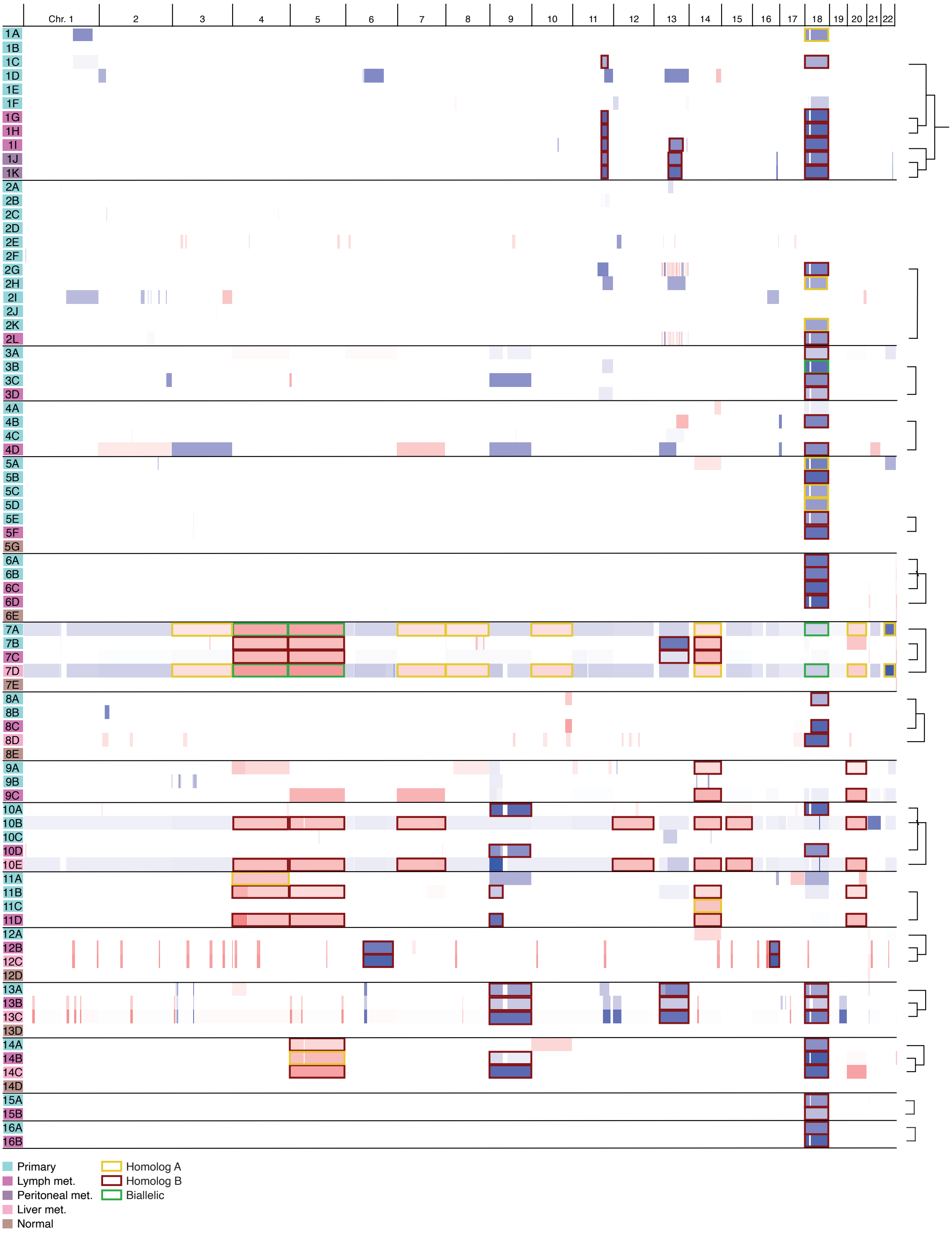


**Copy number alternations and homolog phasing.** Copy number alterations retrieved from Battenberg. The chromosome homolog lost/gained in whole-chromosome and larger recurrent events, determined based on germline SNPs, is indicated by colored borders. SNV-based phylogenies are indicated to the right. Potential subclonal loss of chr18 in 4A, 9C and 11B is not shown in Fig. 1, Fig. 2 or Supplementary Fig. 1 due to low amplitude (logR<0.2).

### Supplementary Figure 3

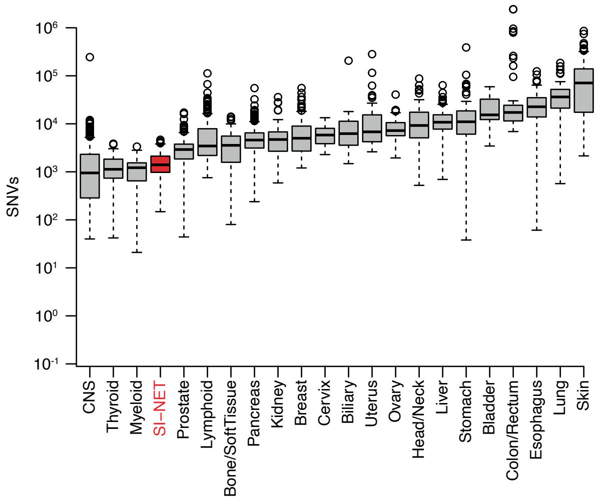


**Genome-wide SNV counts in SI-NET compared to other cancers.** Based on available WGS-derived data from the Pan Cancer Analysis of Whole Genomes (PCAWG) cohort[9].

### Supplementary Figure 4

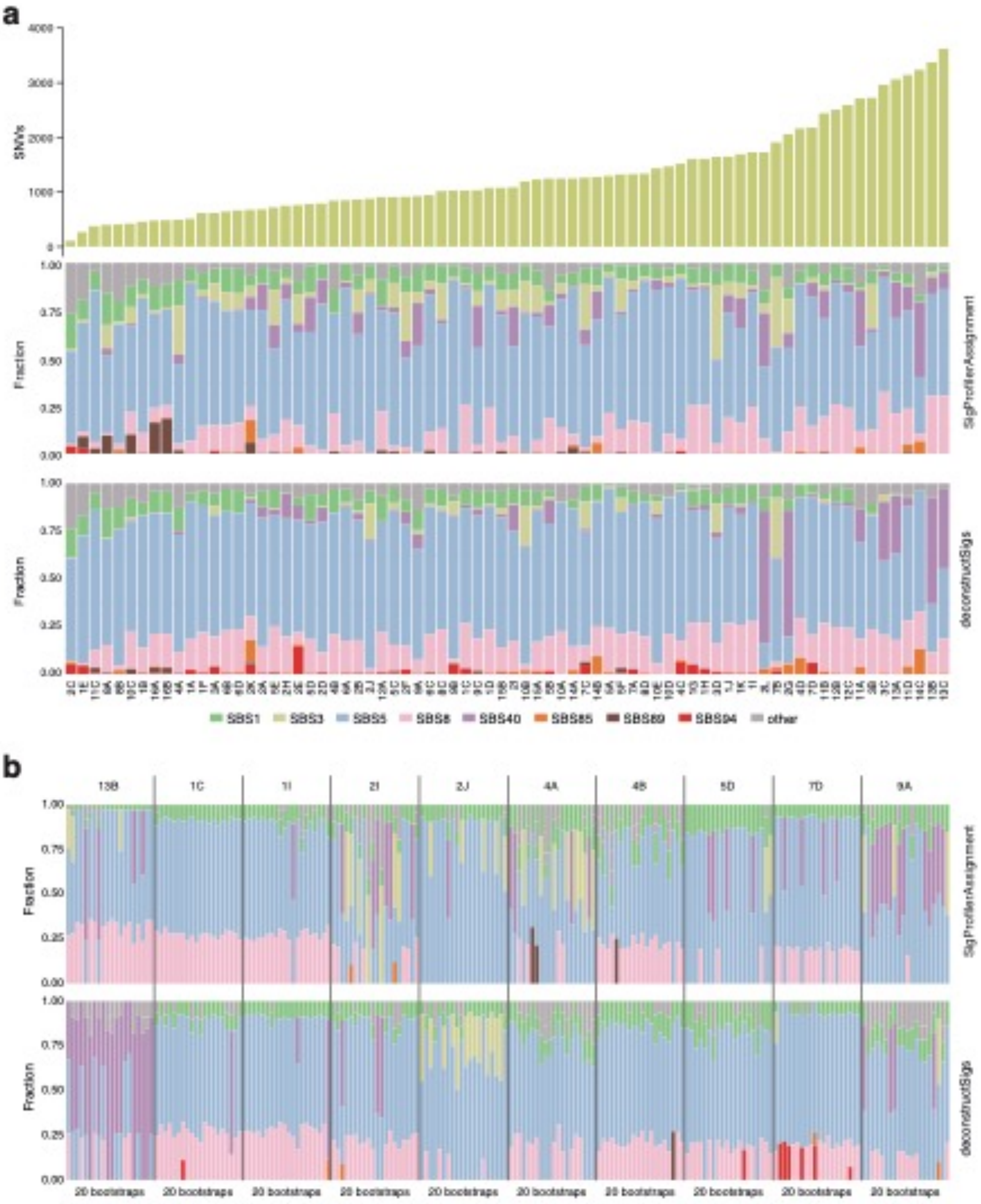


**Mutational signatures in SI-NET are mainly clock-like.** (**a**) Estimated contribution of mutational signatures (COSMIC v3.2) in each tumor sample. The average result from 500 bootstraps from both deconstructSigs and SigProfilerAssignment are shown. Mutation burden is indicated on top. The samples are sorted by increasing mutation burden. (**b**) Individual loadings for the flat/featureless signatures SBS5, SBS40, SBS3 and SBS8 should be interpreted with care. Outcomes from 20 different bootstraps are shown for 10 representative samples, revealing low solution stability with many parameter configurations producing near-equivalent fits.

### Supplementary Figure 5

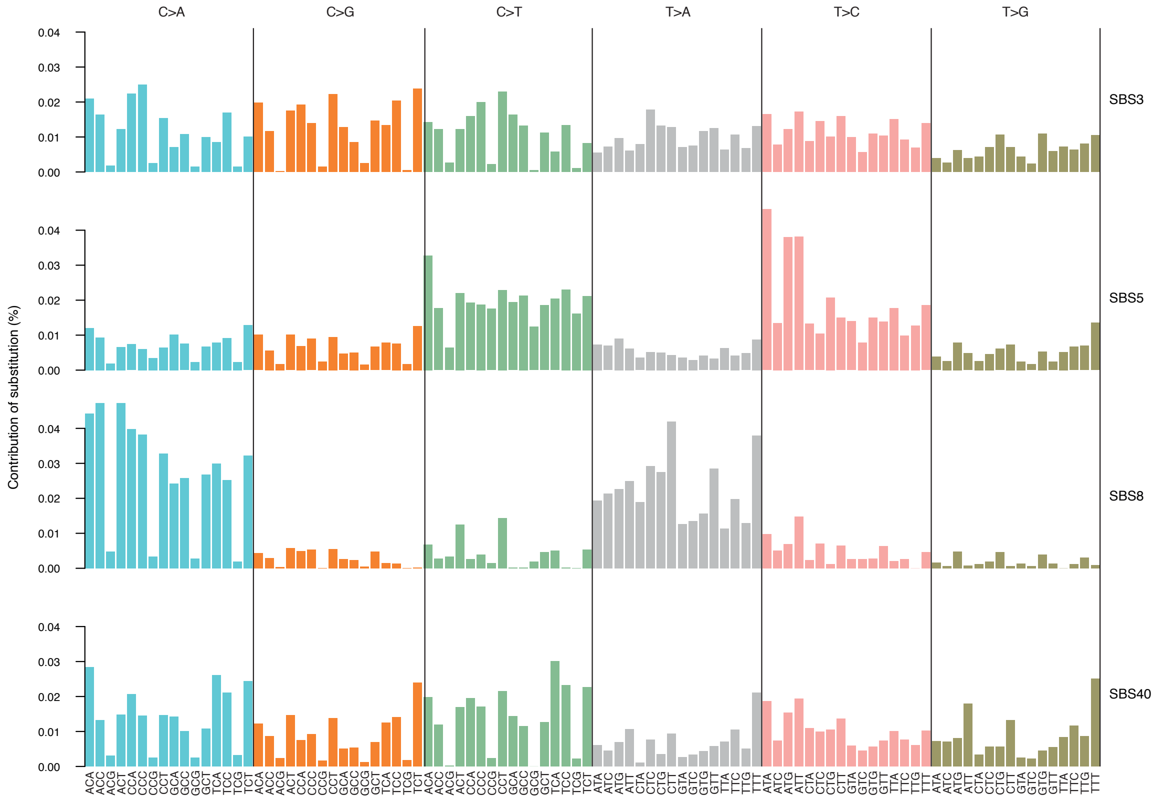


**Key contributing COSMIC signatures in SI-NET have similar trinucleotide spectra.** COSMIC v3.2 trinucleotide spectra for SBS3, SBSB5, SBBS8, SBBS40, which all exhibit relatively unspecific substitution profiles.

### Supplementary Figure 6

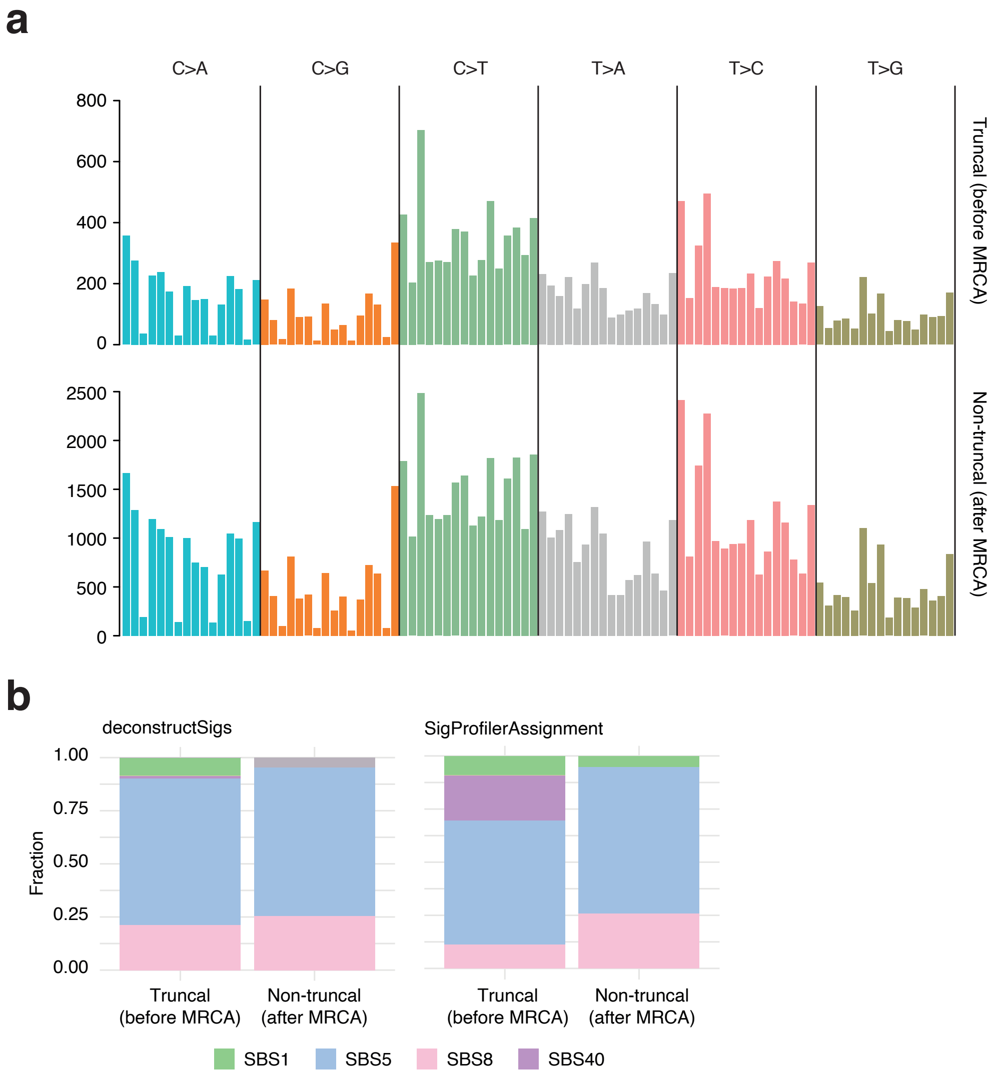


**Mutational signatures are similar in early and late SI-NET mutations.** Mutations in related primary tumors and metastases were separated into early, truncal (shared by metastasis and primary) and later, non-truncal mutations. (**a**) Overall trinucleotide spectra in truncal and non-truncal mutations. (**b**) The average contribution of mutational signatures (COSMIC v3.2) in truncal and non-truncal mutations determined using deconstructSigs and SigProfilerAssignment (average from 500 bootstraps).

### Supplementary Figure 7

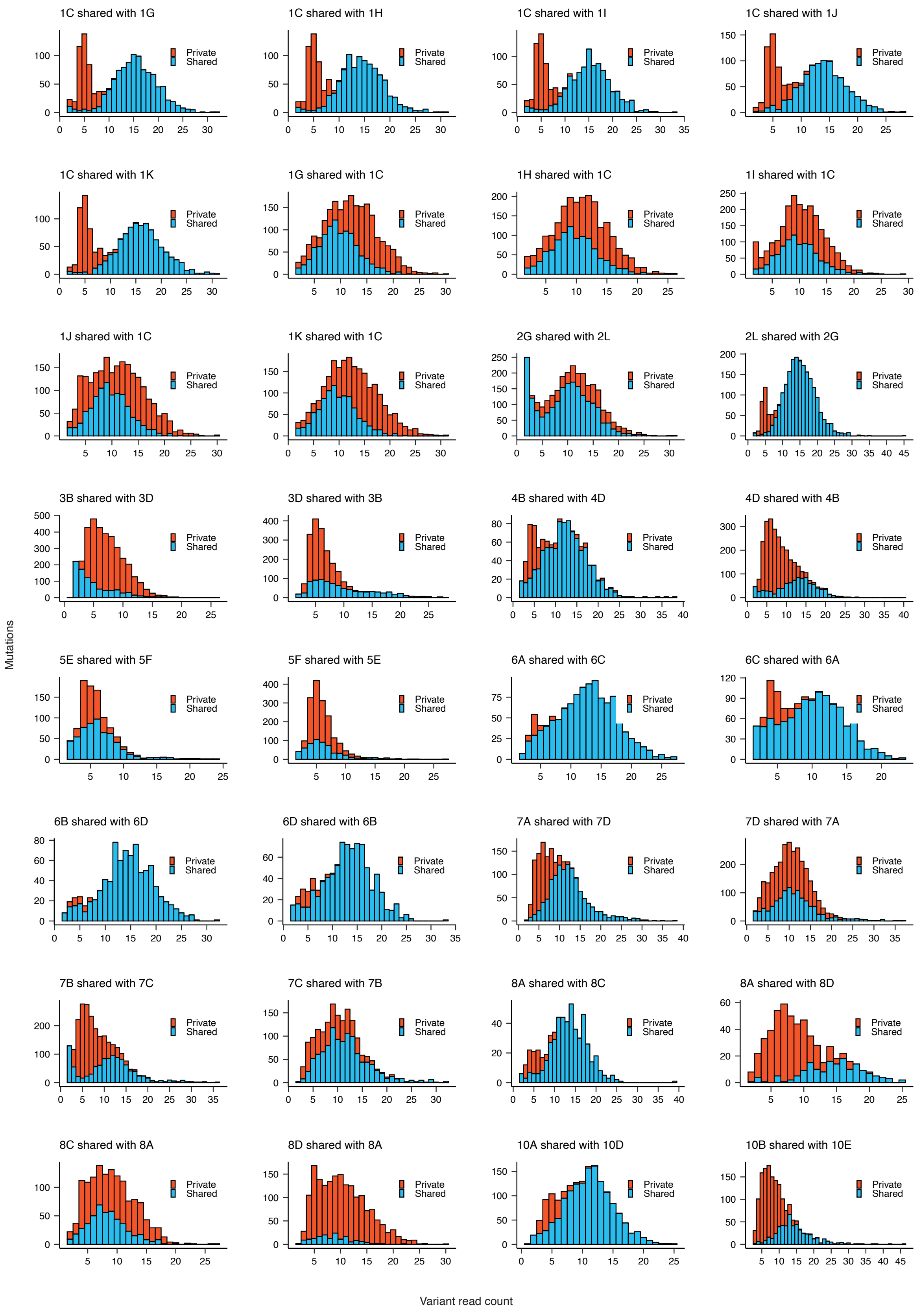


(Continued on the next page)

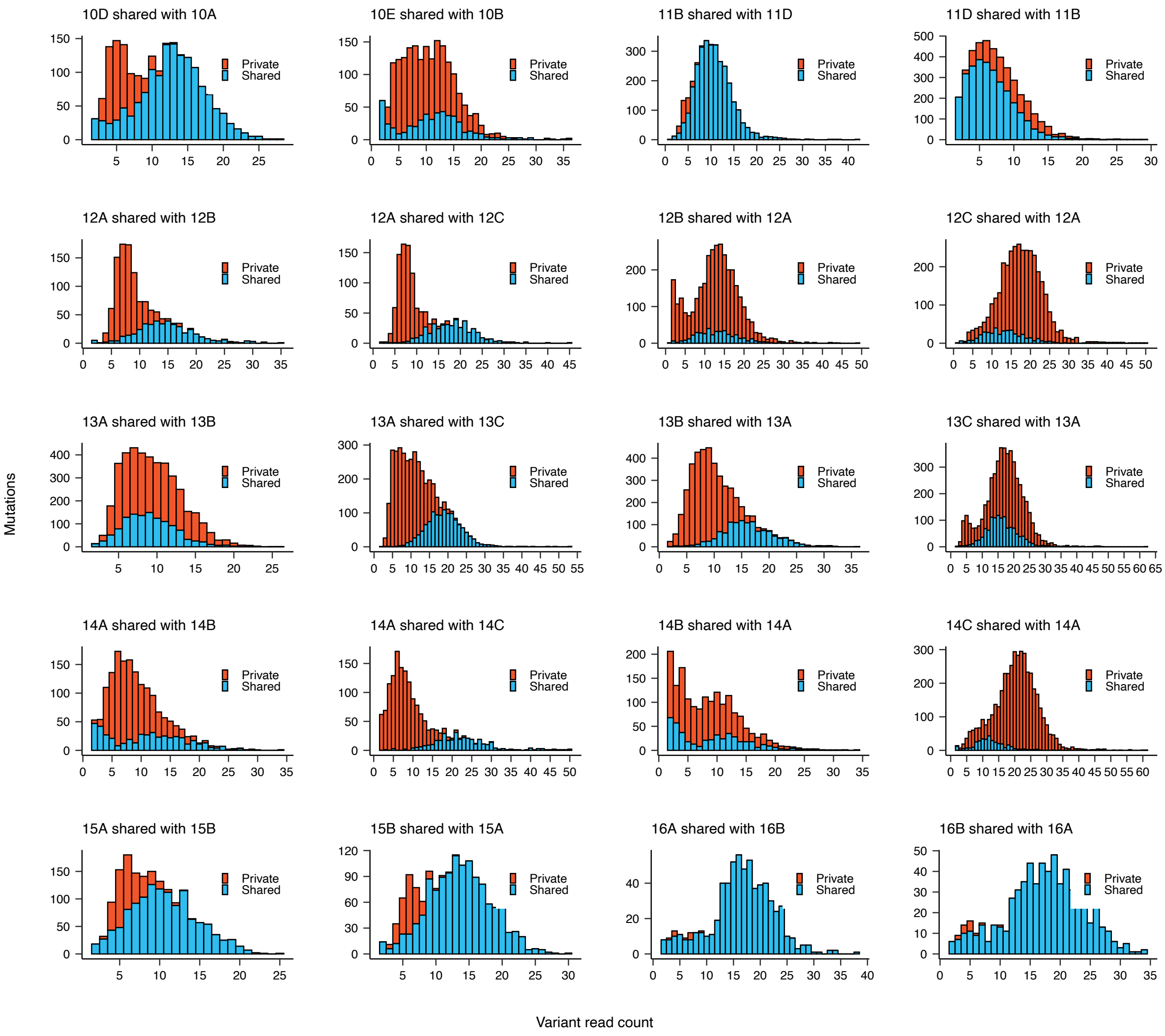


**Histograms of variant allele read counts.** The graphs show the distribution of variant allele read counts for all mutations detected in each tumor shown in main Fig. 2a-b. The histograms are stacked, such that truncal mutations (shared between primary tumors and metastases) and non-truncal mutations are indicated separately. Most distributions fall safely inside the detectable range, implying that drop-outs should be limited.

### Supplementary Note 1

Supplementary Note 1, presented on the following pages, includes detailed information for each patient to support the timelines presented in Figure 2. Overlapping somatic SNVs, phylogenetic relationships, and phasing of CNAs based on heterozygous SNPs are included for all patients. Additional figures supporting timing of WGD and potential driver mutations, are included when needed.

### Patient 1

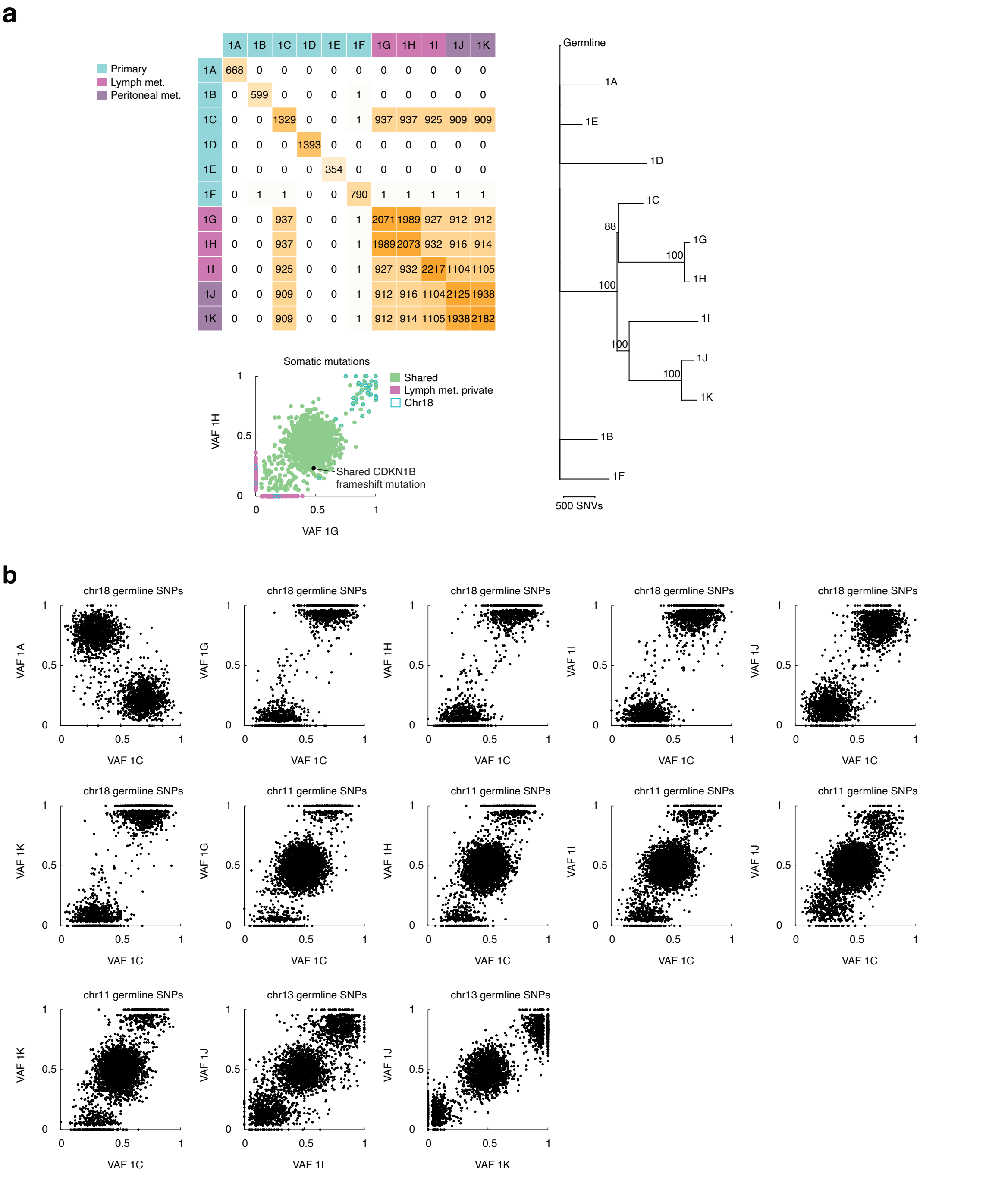


**Patient 1. (a)** Patient 1 has six primary tumours (1A-1F), three lymph node metastases (1G-1I), and two peritoneal metastases (1J-1K). Based on the number of shared mutations, 1C is the seeding primary of 1G-1K. A potential driver in *CDKN1B* is shared only between 1G and 1H, putting this event after 1C:1G-1K MRCA but before 1G:1H MRCA. Potentially, a shared WGD between 1G and 1H can be seen (based on bimodal distribution of shared SNVs). However, k-means clustering could not successfully distinguish the clusters thus not shown in Figure 2. **(b)** Phasing of CNAs based on heterozygous SNPs to support timing. Loss of chr18 and focal loss of chr11 is shared between 1C and its metastases 1G-1K, which puts these events before the 1C:1G-1K MRCA clone. This is further supported by chromosome phasing, where the same homolog is lost in 1C and 1G-1K for both chr11 and chr18. Focal loss of chr13 is shared between 1I, 1J, and 1K and focal loss of chr16 and loss of chr22 are shared between 1J and 1K based on the same reasoning.

### Patient 2

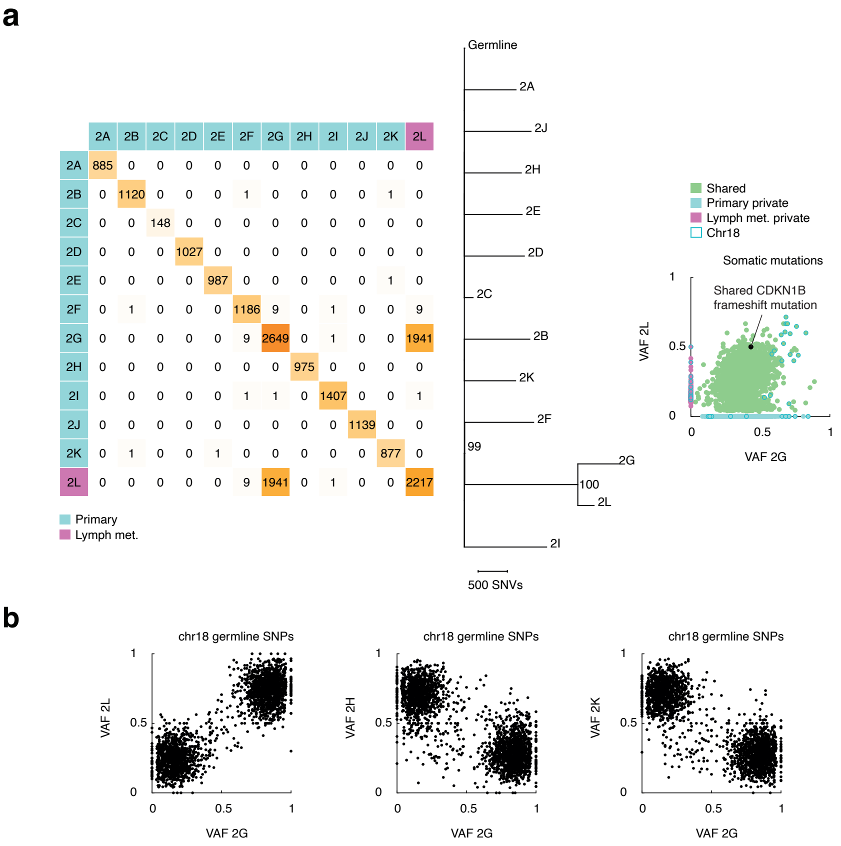


**Patient 2. (a)** Patient 2 has 11 primary tumours (2A-2K) and one lymph node metastasis 2L. Based on the number of shared mutations, 2G is the seeding primary (1941 shared mutations with 2L). Loss of chr18, chromothripsis, and a potential *CDKN1B* driver mutation can be timed before the 2G:2L MRCA clone since these events are shared between the tumours. Focal loss of chr11 in 2G is timed after the 2G:2L MRCA clone since this is private for 2G. **(b)** Phasing of CNAs based on heterozygous SNPs to support timing. Phasing of chr18 loss supports that the event is shared since the same homolog is lost in 2G and 2L.

### Patient 3

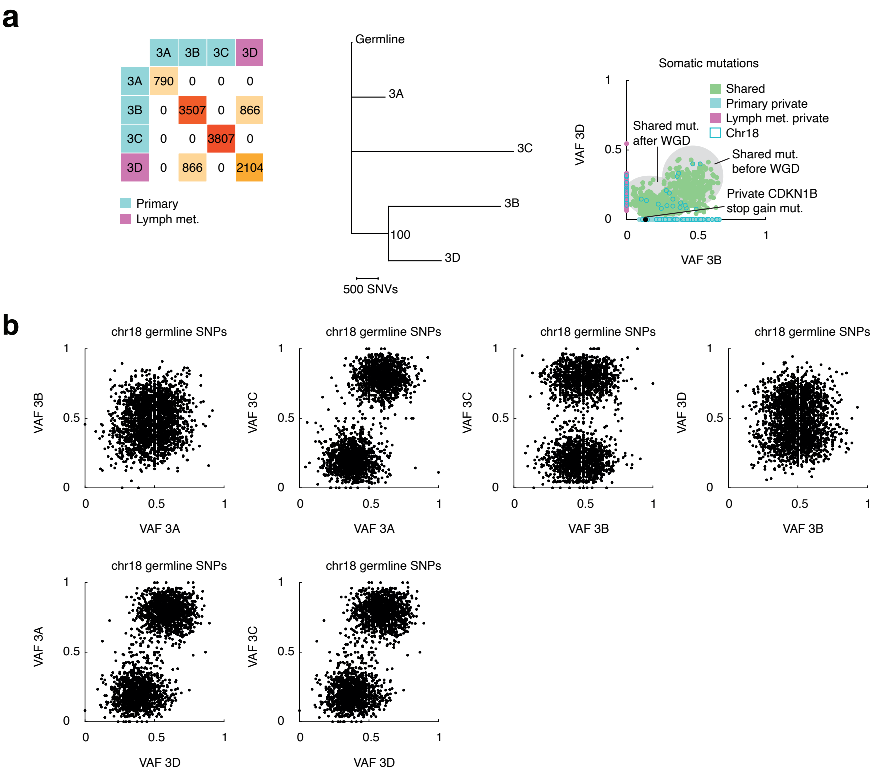


**Patient 3. (a)** Patient 3 has three primary tumours (3A-3C) and one lymph node metastasis 3D. Based on the number of shared mutations, 3B is the seeding metastasis (866 shared mutations with lymph node met. 3D). Based on VAF distribution of shared mutations early WGD before 3B:3D MRCA is likely (271 mutations before, 564 mutations after based on k-means clustering). Potential driver mutation in *CDKN1B* is timed after 3B:3D MRCA clone since this is a private event in 3B. **(b)** Phasing of CNAs based on heterozygous SNPs to support timing. Based on chromosome phasing loss of chr18 is biallelic in 3B thus must have occurred after WGD. In 3D, one likely explanation is that 1 out of 4 chromosome copies are lost after WGD, based on chromosome phasing (clusters slightly separated from 0.5). This is also supported by the difference in CNA amplitude between 3B and 3D (see Supplementary Figure 2) where the amplitude in 3B is lower than 3D supporting loss of 2 out 4 respectively 1 out of 4 chromosomes copies after WGD. The amplitude difference and cluster separation could also be explained by low purity in 3D thus 2 chromosome copies are lost (one of each homolog) but affected by copy number neutral normal tissue. Loss of 1 of the chromosome copies after WGD are most likely a shared event between 3B and 3D putting this before the 3B:3D MRCA clone.

### Patient 4

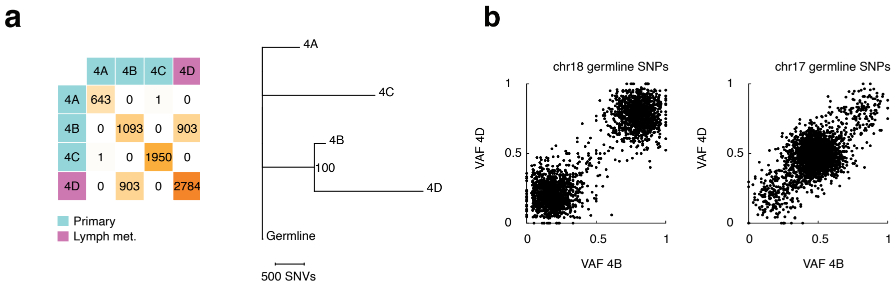


**Patient 4. (a)** Patient 4 has three primary tumours (4A-4C) and one lymph node metastasis 4D. Based on the number of shared mutations, 4B is the seeding primary (903 shared mutation with lymph node metastasis 4D). **(b)** Phasing of CNAs based on heterozygous SNPs to support timing. Loss of chr18 and focal loss of chr17 can be timed before the 4B:4D MRCA clone since these events are shared between the tumours and the same homologs are lost in both tumours based on chromosome phasing. Other chromosome gains and losses in 4B and 4D are timed after the 4B:4D MRCA clone since these are private for 4B respective 4D.

### Patient 5

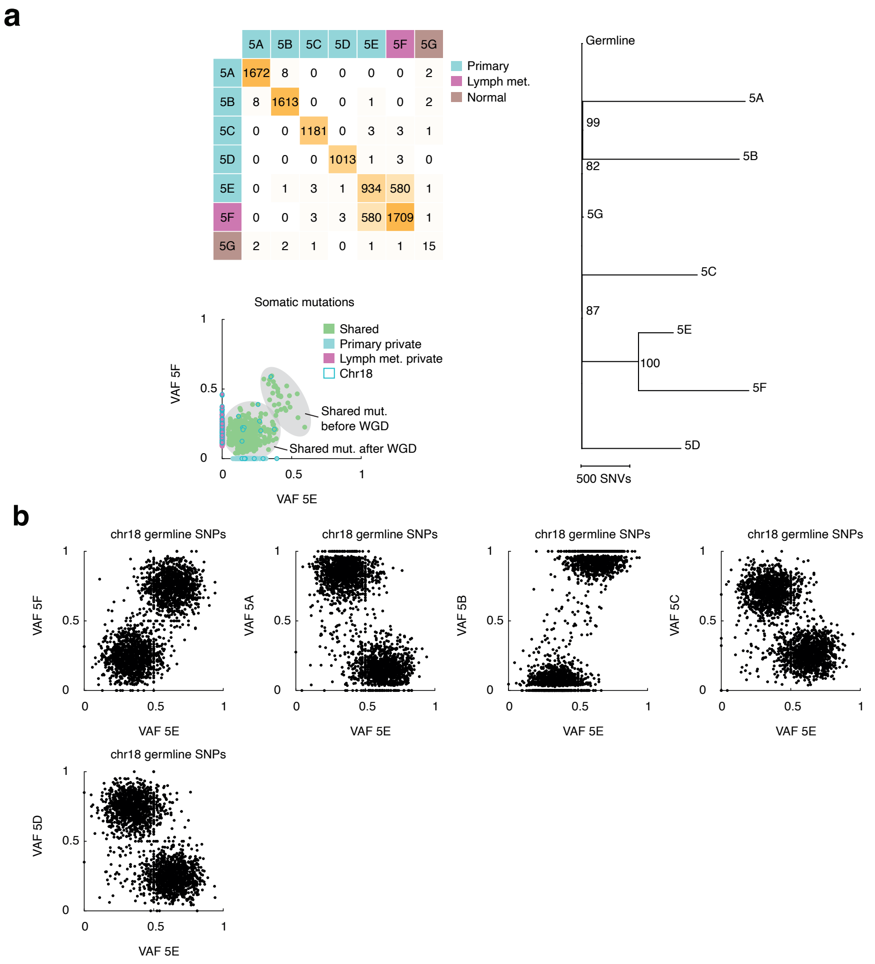


**Patient 5. (a)** Patient 5 has five primary tumours (5A-5E), one lymph node metastasis 5F and one normal sample 5G. Based on the number of shared mutations, 5E is the seeding primary (580 shared mutation shared with lymph node met. 5F). WGD is timed before MRCA. The bimodal distribution of shared mutations indicates that the event is shared (39 mutations before, 482 mutations after based on k-means clustering). It is likely that WGD occurred before loss of chr18 since the private mutations have a slightly higher VAF then CN neutral chromosomes, as expected when there are 3 instead of 4 chromosome copies. **(b)** Phasing of CNAs based on heterozygous SNPs to support timing. Loss of chr18 is timed before the 5E:5F MRCA clone since this event is shared, and the same homolog is lost in both samples.

### Patient 6

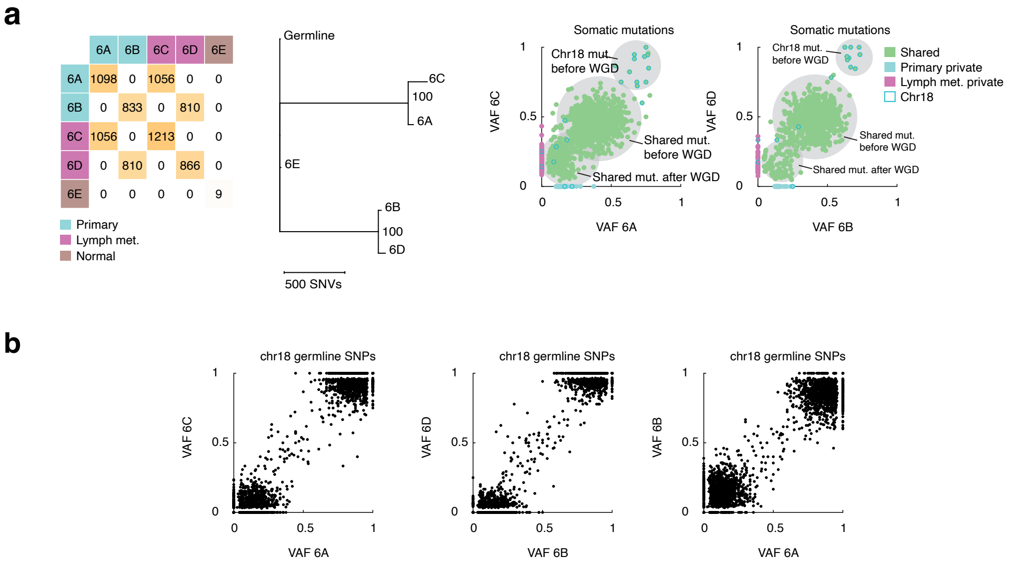


**Patient 6. (a)** Patient 6 has two primary tumours (6A-6B), two lymph node metastases (6C-6D) and one normal sample 6E. Based on the number of shared mutations 6A is the seeding primary to 6C (1056 shared mutation) and 6B is the seeding primary to 6D (810 shared mutations). Loss of chr18, followed by WGD can be timed before MRCA clone in both pairs (6A:6C and 6B:6D). In both cases WGD is based on the bimodal VAF distribution of shared mutations (779 before and 258 after in 6A:6C and 665 before and 124 after in 6B:6D). **(b)** Phasing of CNAs based on heterozygous SNPs to support timing. Chr18 loss is shared between the tumours pairs and same homolog is lost in both pairs, putting these events before the 6A:6C respectively 6B:6D MRCA clone.

### Patient 7

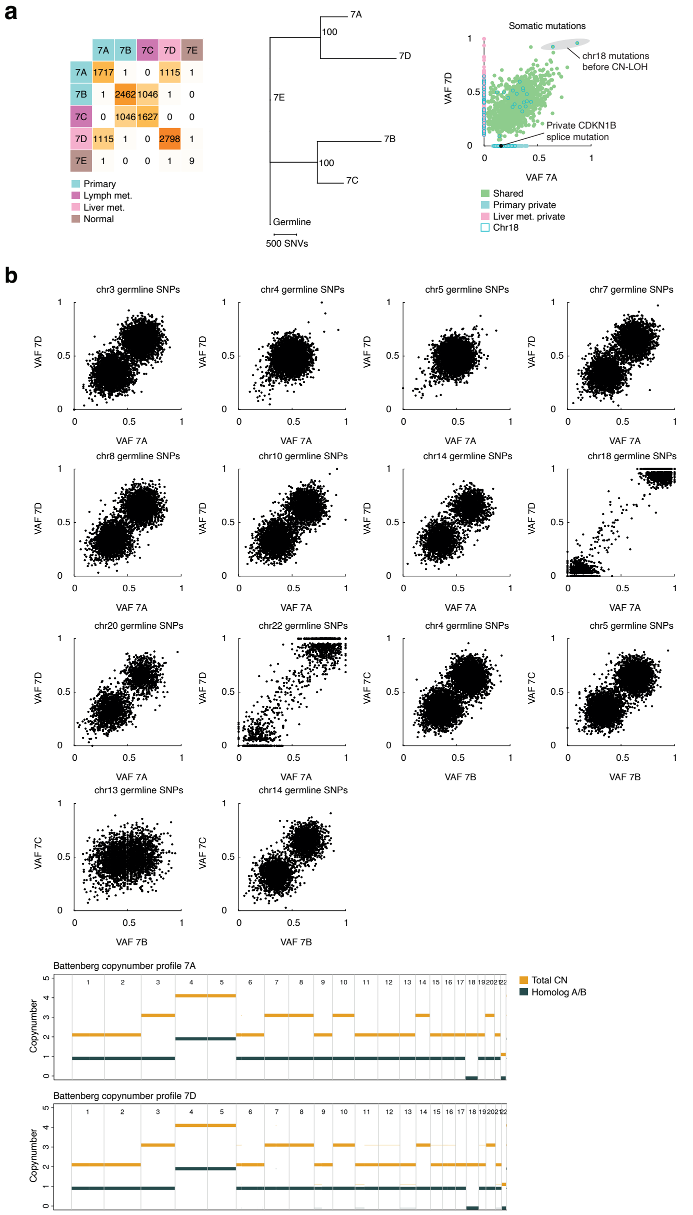


**Patient 7. (a)** Patient 7 has two primary tumours (7A-7B), one lymph node metastasis 7C, one liver metastasis 7D and one normal sample 7E. Based on the number of shared mutations, 7A is the seeding primary to 7D (1115 shared mutations) and 7B is the seeding primary to 7C (1046 shared mutations). In 7A and 7D CN-LOH of chr18 can med timed before 7A:7D MRCA clone. Based on results from Battenberg (panel c) two copies of chr18 are present, both from the same homolog. This event is timed very early since shared mutation before CN-LOH should have a high VAF ~1. There are two shared high VAF mutations on chr18. Early mutations on the lost homolog cannot be seen in the data, but if the same mutation frequency acts on both homologs, 4 mutations can be assumed on chr18 before CN-LOH. Potential *CDKN1B* driver mutation in 7A is timed after 7A:7D MRCA clone, since this event is unique for 7A. **(b)** Phasing of CNAs based on heterozygous SNPs to support timing. Loss of chr22 and gain of chr3,4,5,7,8,10,14, and 20 can also be timed before the 7A:7D MRCA clone, all supported by chromosome phasing. Loss of chr22 and gain of chr 3,7,8,10,14, and 20 are supported by chromosome phasing where the same homolog is lost or gained in both samples. Gain of chr4 and chr5 is biallelic and one of each homolog is gained in both samples. In 7B and 7C, loss of chr13 and gain of chr4,5,14 can be timed before the 7B:7C. This is supported by chromosome phasing. Gain of chr20 and focal gains in

chr3,8 are timed after the 7B:7C MRCA clone since these events are private to 7B. **(c)** Results from Battenberg supporting CN-LOH of chr18 in 7A and 7D.

### Patient 8

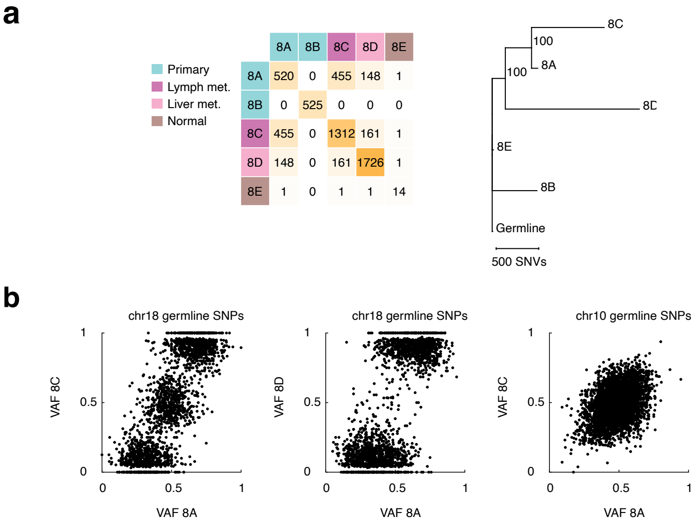


**Patient 8. (a)** Patient 8 has two primary tumours (8A-8B), one lymph node metastasis 8C, one liver metastasis 8D and a normal sample 8E. Based on the number of shared mutations, 8A is the seeding primary to 8C and 8D. **(b)** Phasing of CNAs based on heterozygous SNPs to support timing. 18q is shared between all tumours, and the same homolog is lost in all three samples, and was therefore assumed to be a truncal event, although potentially there could be two separate, later, events (whole-chromosome loss in 8D and 18q loss shared by 8A and 8C). Gain of chr10 can be timed after 8A:8B-8C MRCA clone but before 8A:8C MRCA clone since this is shared between 8A and 8C. Gain of chr9 and chr20 are timed after 8A:8B-8C since these are private to 8D.

### Patient 9

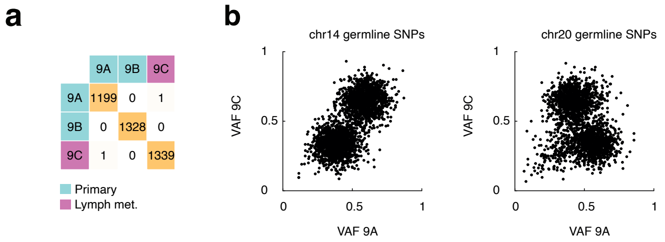


**Patient 9. (a)** Patient 9 has two primary tumours (9A-9B) and one lymph node metastasis 9C. None of the primary tumours is the seeding primary to 9C. **(b)** Phasing of CNAs based on heterozygous SNPs. The same homolog of chr14 and different homologs of chr20 was lost in 9A and 9C.

### Patient 10

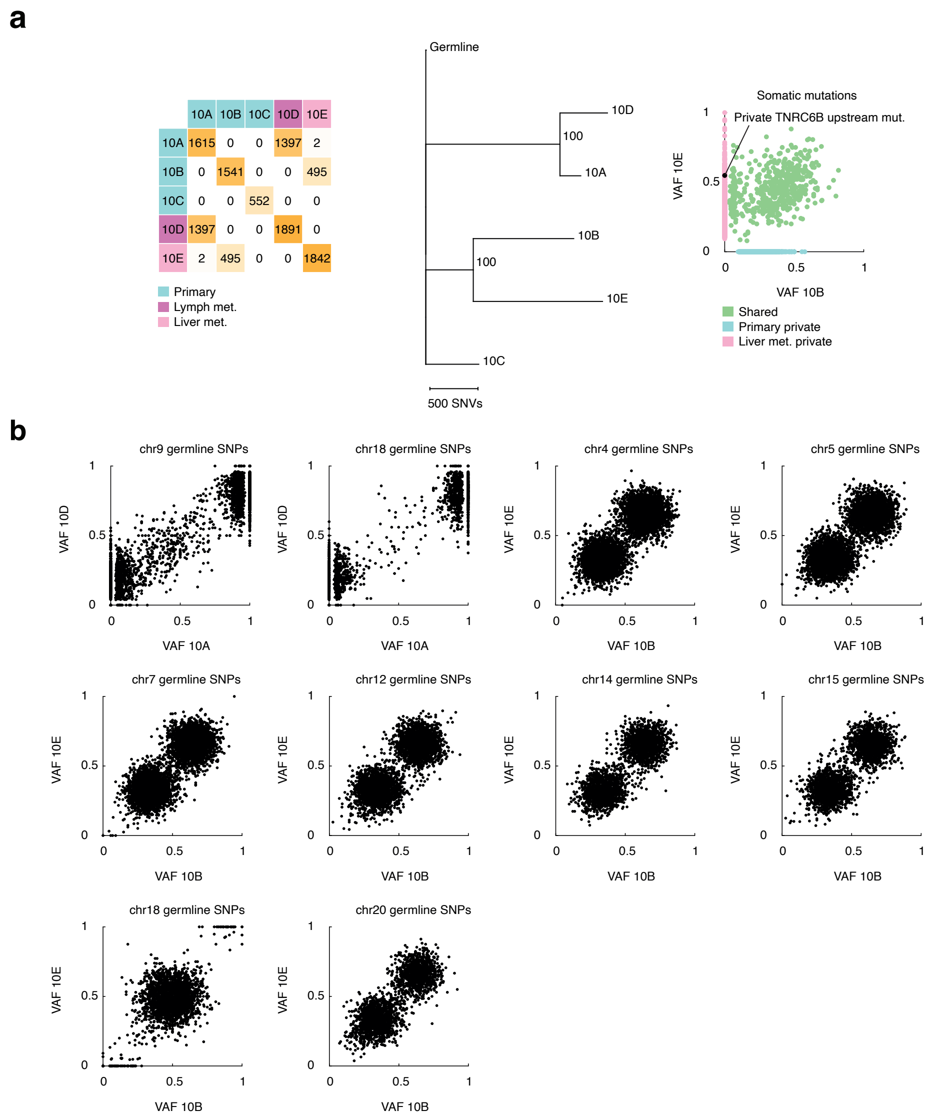


| | 10A | 10B | 10C | 10D | 10E |
|---|---|---|---|---|---|
| 10A | 1615 | 0 | 0 | 1397 | 2 |
| 10B | 0 | 1541 | 0 | 0 | 495 |
| 10C | 0 | 0 | 552 | 0 | 0 |
| 10D | 1397 | 0 | 0 | 1891 | 0 |
| 10E | 2 | 495 | 0 | 0 | 1842 |

**Patient 10. (a)** Patient 10 has three primary tumours (10A-10C), one lymph node metastasis 10D, and one liver metastasis 10E. 10A is the seeding primary to 10D (1397 shared mutations) and 10B is the seeding primary to 10E (495 shared mutations). Potential driver in *TNRC6B* is timed after the 10B:10E MRCA clone since this event is private to 10E. **(b)** Phasing of CNAs based on heterozygous SNPs to support timing. 10A and 10D share loss of chr18 and chr9 which puts these events before 10A:10D MRCA clone. This is supported by chromosome phasing. 10B and 10C share a focal loss in chr18 and gains in chr4,5,7,12,14,15, and 20 and all these events are therefore timed before 10B:10C MRCA clone. This is supported by chromosome phasing. Additional chromosome losses are timed after the 10B:10E MRCA clone since these events are private to 10B respectively 10E.

### Patient 11

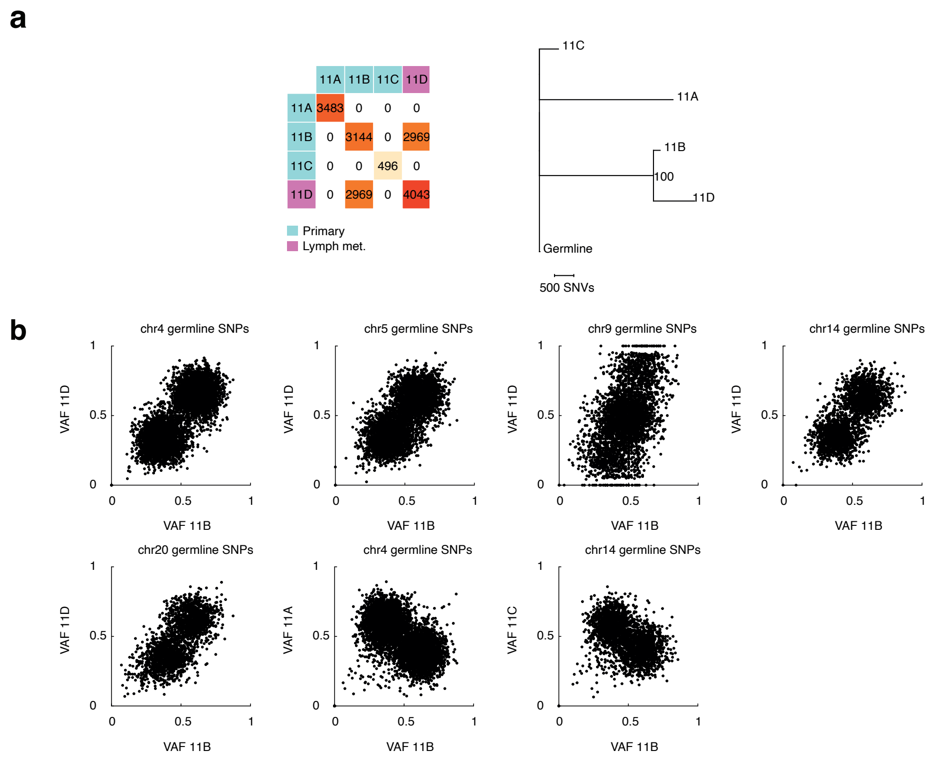


**Patient 11. (a)** Patient 11 has three primary tumours 11A-11C and one lymph node metastasis 11D. Based on the number of shared mutations, 11B is the seeding primary (2969 shared mutations with 11D). **(b)** Phasing of CNAs based on heterozygous SNPs to support timing. Gain of chr4,5,14,20 and loss of chr9p can be timed before 11B:11D MRCA clone since these events are shared. This is also supported by chromosome phasing.

### Patient 12

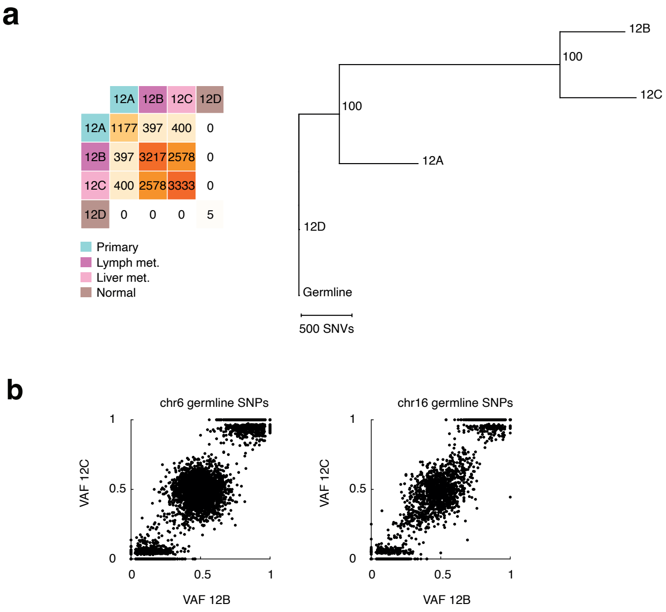


**Patient 12. (a)** Patient 12 (unifocal) has one primary tumour 12A, one lymph node metastasis 12B, one liver metastasis 12C, and one normal sample. No events can be timed before the 12A:12B-12C MRCA clone. **(b)** Phasing of CNAs based on heterozygous SNPs to support timing. Focal gains and losses of multiple chromosomes can be timed after 12A:12B-12C MRCA clone but before 12B:12C MRCA clone, since these events are shared between 12B and 12C. Losses in chr6 and chr16 is supported by chromosome phasing. Other focal events were too small to perform chromosome phasing. Gain of chr14 is timed after 12A:12B-12C MRCA clone and is private to 12A.

### Patient 13

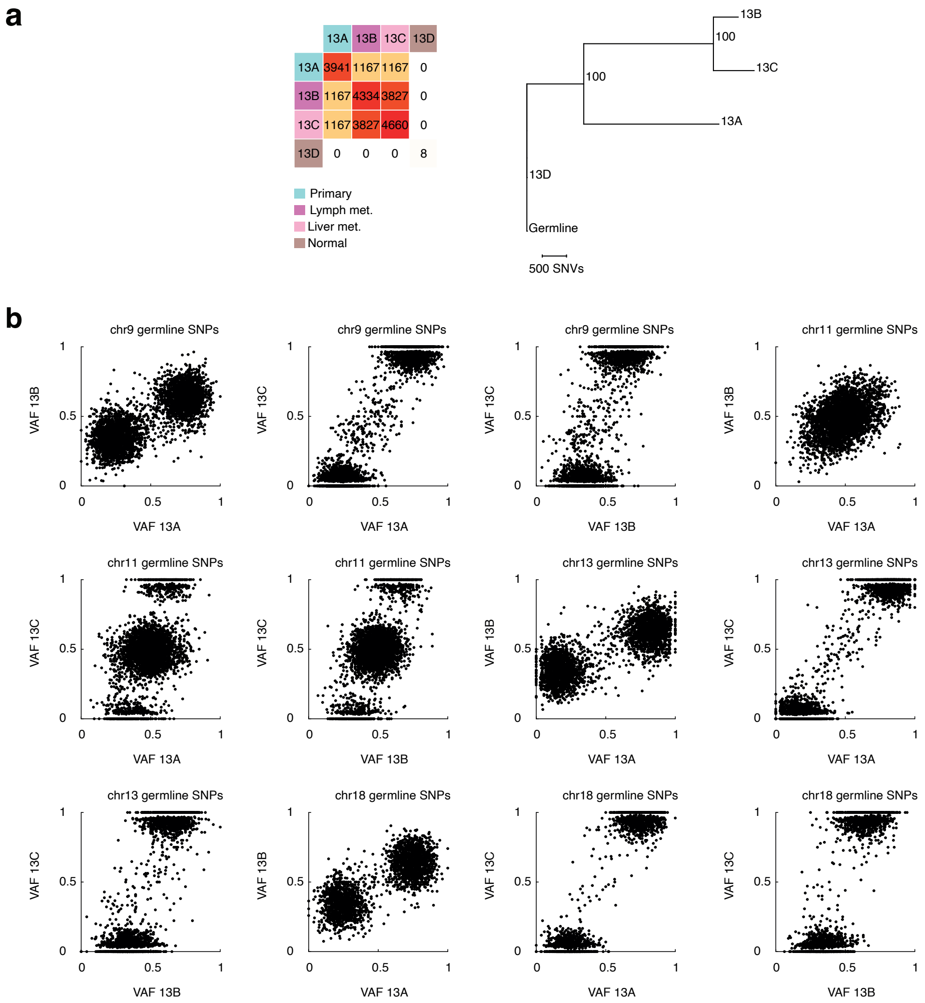


**Patient 13. (a)** Patient 13 (unifocal) has one primary tumour 13A, one lymph node metastasis 13B, one liver metastasis 13C and on normal sample. **(b)** Phasing of CNAs based on heterozygous SNPs to support timing. Loss of chr9, chr13, and chr18 is shared between all tumours and is timed before the 13A:13B-13C MRCA clone. This is also supported by chromosome phasing. Focal loss of chr3 and chr6 is shared between all samples and is therefore timed before the 13A:13B-13C MRCA clone. Since the losses are very small, no phasing was performed, but it is unlikely that these are separate events. Focal losses and gains of multiple chromosomes are likely shared between 13B and 13C and is therefore timed after 13A:13B-13C MRCA clone but before 13B:13C MRCA clone. Focal loss of chr11 is private to 13A.

### Patient 14

**a**

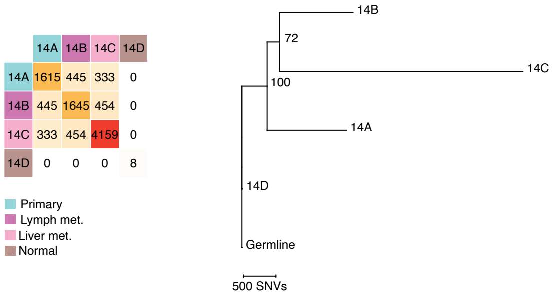


**b**

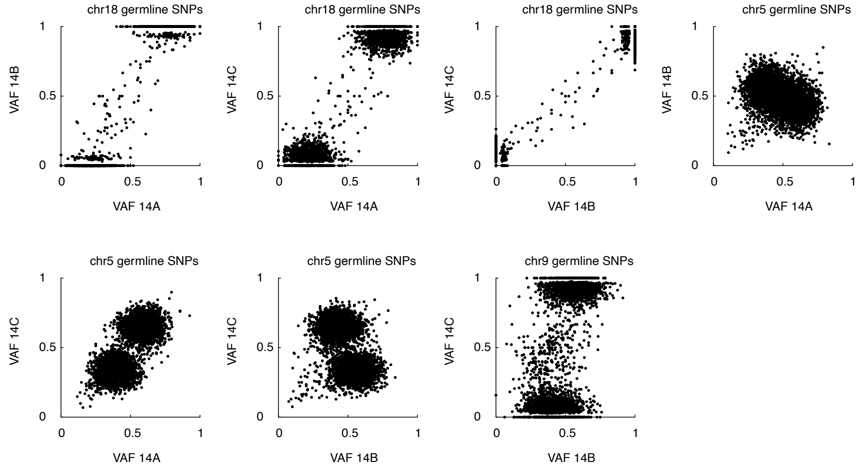


**Patient 14. (a)** Patient 14 (unifocal) has one primary tumour 14A, one lymph node metastasis 14B, one liver metastasis 14C and a normal sample. **(b)** Phasing of CNAs based on heterozygous SNPs to support timing. Loss of chr18 is shared between all tumours and is timed before the 14A:14B-14C MRCA clone. This is also supported by chromosome phasing which shows that the same homolog is lost in all three tumours. The same reasoning is used to time loss of chr9 after 14A:14B-14C but before the 14B:14C MRCA clone. Chromosome phasing of chr5 indicates that the same homolog is gained in 14A and 14C, while the other homolog is gained in 14B. Based on this, it cannot be excluded that these are three separate events. Gain of chr10 and chr20 are private to 14A respectively 14C.

### Patient 15

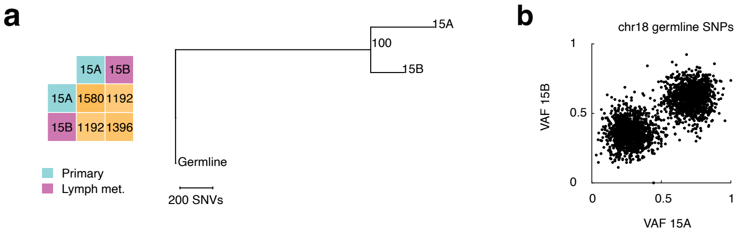


**Patient 15. (a)** Patient 15 (unifocal) has one primary tumour 15A and one lymph node metastasis 15B that share 1192 mutations. **(b)** Phasing of CNAs based on heterozygous SNPs to support timing. Loss of chr18 is a shared event between 15A and 15B and is timed before 15A:15B MRCA clone. This is also supported by homolog phasing which shows that the same homolog is lost in both tumours.

### Patient 16

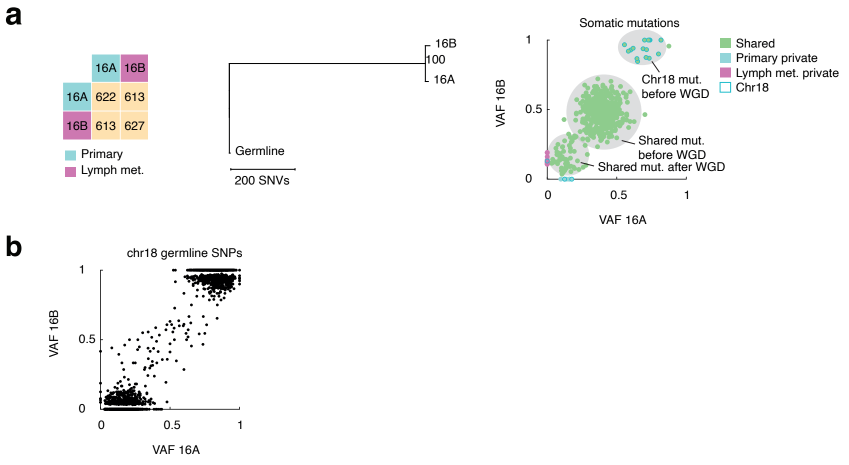


**Patient 16. (a)** Patient 16 (unifocal) has one primary tumour 16A and one lymph node metastasis 16B that share 613 mutations. WGD is also timed before MRCA clone, but after chr18 loss. The bimodal distribution of shared mutations indicates that the event is shared (463 mutations before, 70 mutations after based on k-means clustering). **(b)** Phasing of CNAs based on heterozygous SNPs to support timing. Loss of chr18 is a shared event between 16A and 16B and is timed before 16A:16B MRCA clone. This is also supported by chromosome phasing which shows that the same homolog is lost in both tumours.

### Supplementary Note 2

Supplementary Note 2 contains result from MutationTimeR for primary tumor-metastases pairs which have shared copy number gains (7A:7D, 7B:7C, 10B:10E, 11B:11D, 14A:14B-14C). Samples with only copy number losses were not considered since these are not possible to time based on SNVs.

### Patient 7

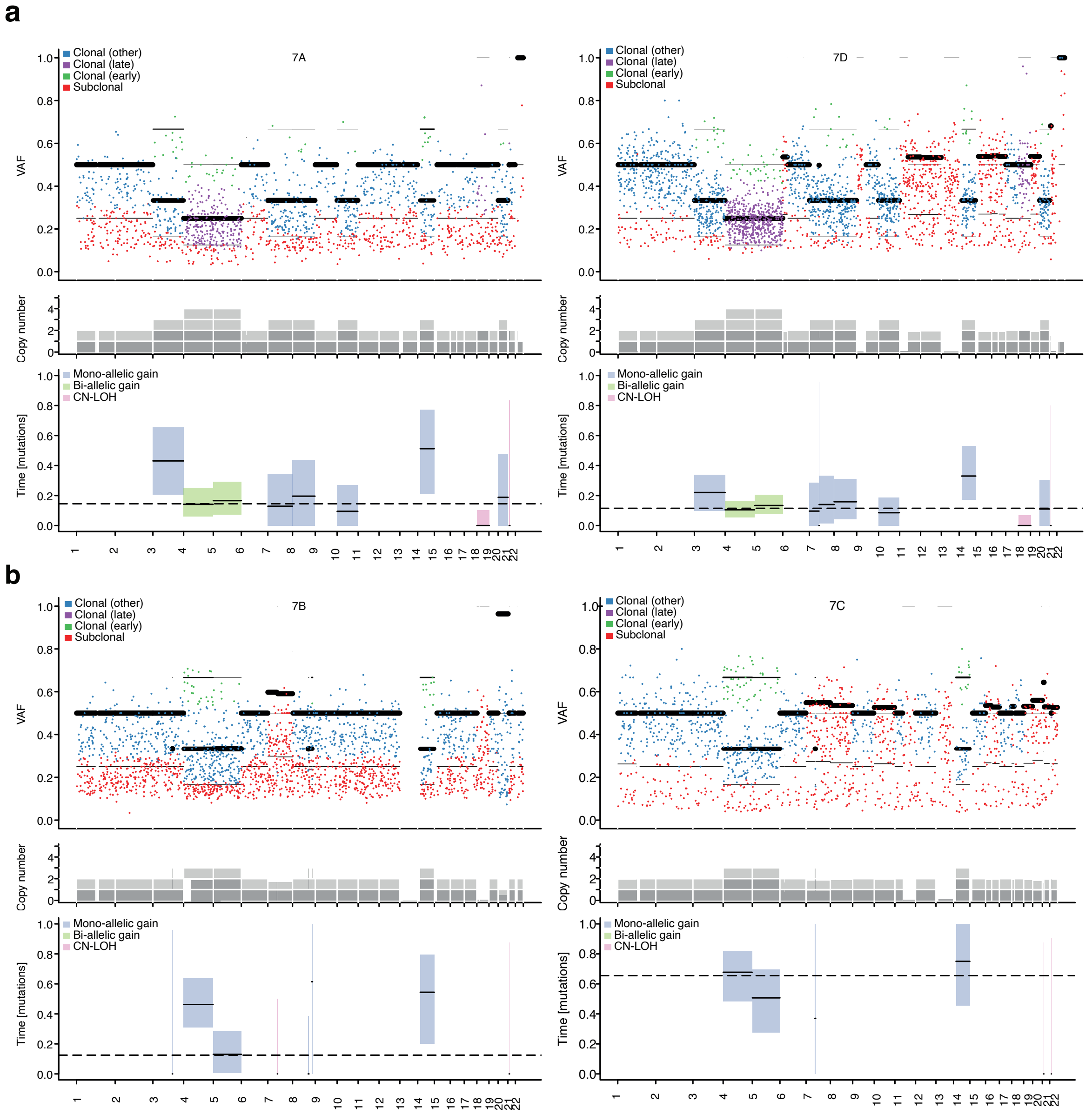


**Patient 7. (a)** Results from MutationTimeR support early CN-LOH shared between 7A and 7D. Biallelic gain in chr4 and chr5 and monoallelic gain in chr3,7,8,10,14,20 at similar timepoints in 7A and 7D are also supported. **(b)** Gains in chr4,5,14 at similar timepoints in 7B and 7C are supported by MutationTimeR.

### Patient 10

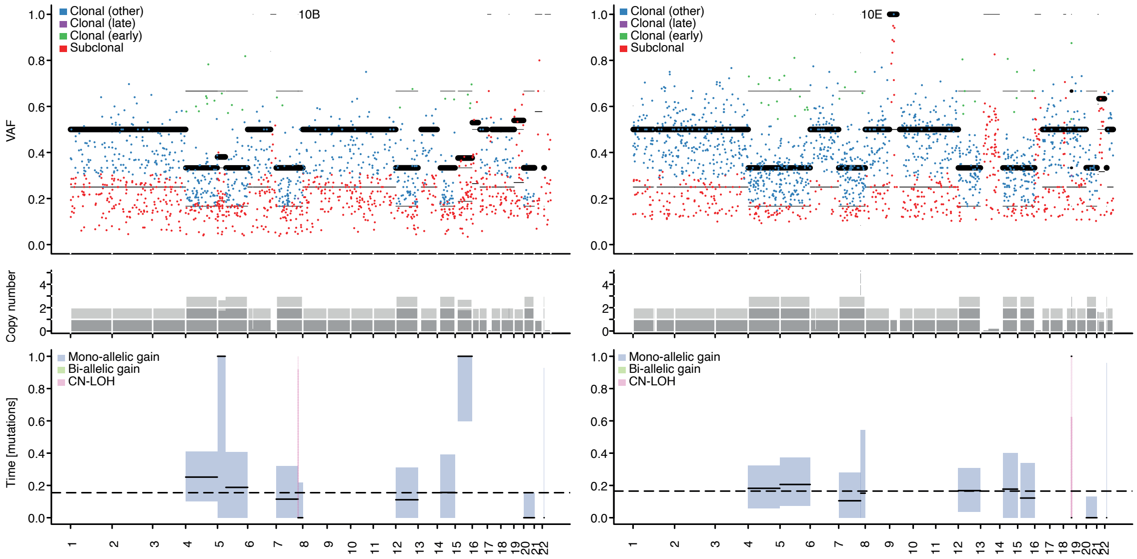


**Patient 10.** Gains in chr4,5,7,12,14,20 at similar timepoints in 10B and 10D are supported by MutationTimeR. Gain of chr15 is timed differently in 10B and 10E. However, this is supported to be a shared event based on chromosome phasing (Supplementary Note 1).

### Patient 11

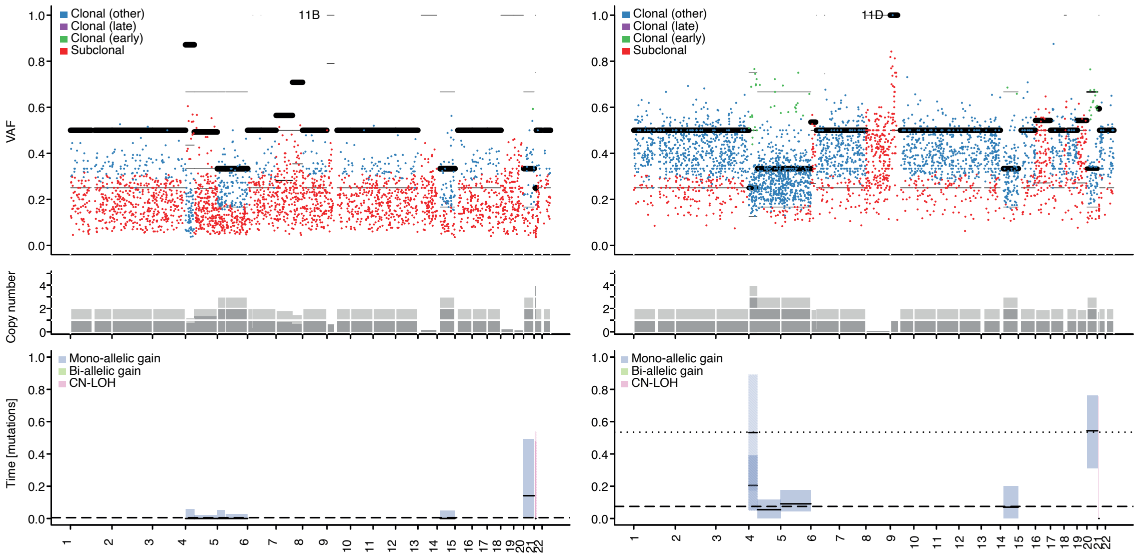


**Patient 11.** Gains in chr4,5,14,20 at similar timepoints in 11B and 11D are supported by MutationTimeR.

### Patient 14

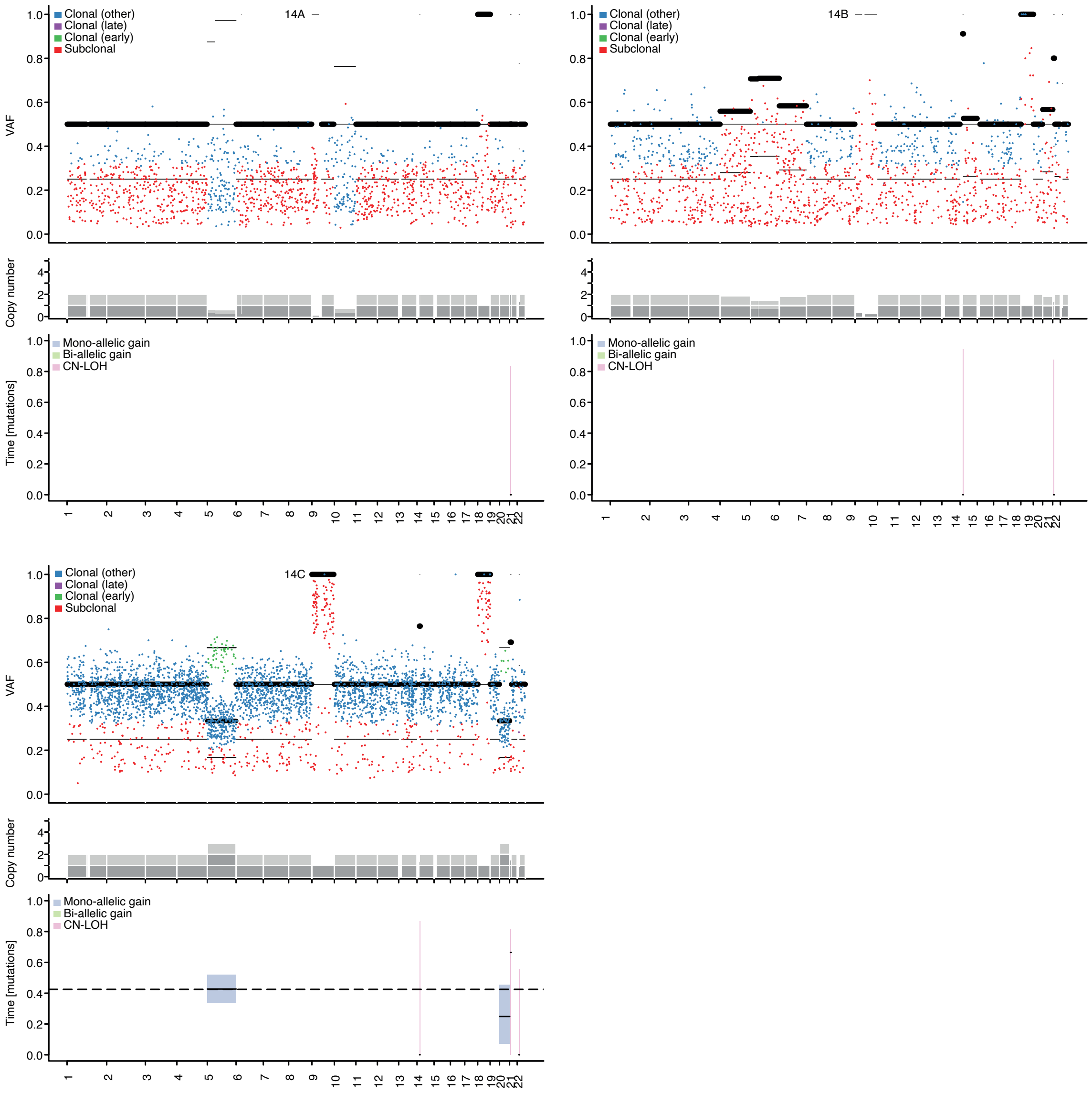


**Patient 14.** No gains could be timed in 14A and 14B. In 14C, gains in chr5 and chr20 could be timed. Late timing of chr5 (after private chr20 gain) supports that this is a private event in 14C, as indicated by chromosome phasing (Supplementary Note 1).
